# ESCRT-dependent STING degradation curtails steady-state and cGAMP-induced signaling

**DOI:** 10.1101/2022.09.22.509044

**Authors:** Matteo Gentili, Bingxu Liu, Malvina Papanastasiou, Deborah Dele-Oni, Marc A Schwartz, Rebecca J. Carlson, Aziz Al’Khafaji, Karsten Krug, Adam Brown, John G Doench, Steven A Carr, Nir Hacohen

**Author notes:** Correspondence (N.H.).

## Abstract

STING is an intracellular sensor of cyclic di-nucleotides involved in response to pathogen- or self-derived DNA that induces protective immunity, or if dysregulated, autoimmunity. STING trafficking is tightly linked to its activity. We aimed to systematically characterize genes regulating STING trafficking and to define their impact on STING responses. Based on proximity-ligation proteomics and genetic screens, an ESCRT complex containing HGS, VPS37A and UBAP1 was found to be required for STING degradation and signaling shutdown. Analogous to phosphorylated STING creating a platform for IRF3 recruitment, oligomerization-driven STING ubiquitination by UBE2N formed a platform for ESCRT recruitment at the endosome, responsible for STING signaling shutdown. A UBAP1 mutant that underlies human spastic paraplegia and disrupts ESCRT function led to STING-dependent type I IFN responses at the steady-state, defining ESCRT as a homeostatic regulator of STING signaling.

## Introduction

Intracellular DNA is a potent activator of innate immune responses via Stimulator of Interferon Genes (STING)^1^, which acts as an adaptor for cyclic GMP-AMP (cGAMP) after its generation by the DNA sensor cyclic GMP-AMP (cGAMP) Synthase (cGAS)^2–5^. In addition to the eukaryotic 2’3’-linked cGAMP, STING is also a sensor for 3’3’-linked cyclic-dinucleotides (CDNs) of bacterial origin^6, 7^. STING binding to its ligands induces type I interferon (IFN) and NF-κB responses in addition to autophagy, an ancestral and conserved activity important for clearance of intracellular pathogens^8^. Activation of innate immune pathways by STING is highly conserved from metazoans to bacteria^9, 10^. Along with anti-pathogen responses, the cGAS/STING axis is essential for antitumor immune responses, immune checkpoint therapy, development of autoimmune diseases and induction of cellular senescence^11^.

Homodimeric STING is localized at the endoplasmic reticulum (ER)^12^ and undergoes a cGAMP-dependent conformational switch that triggers its exit from the ER and trafficking to the Golgi^12, 13^. STING palmitoylation at the Golgi is required for STING clustering and activation of type I IFN responses via Tank Binding Kinase 1 (TBK1) at the Trans-Golgi Network (TGN)^14^. After TBK1 phosphorylates STING at residue S366, phospho-STING forms a platform at the carboxy-terminal tail (CTT) for recruitment of IRF3^15^. IRF3 is then phosphorylated by TBK1, homodimerizes and translocates to the nucleus resulting in activation of a type I IFN response. In addition to type I IFN induction, upon CDN ligation, STING activates NF-κB and induces autophagy independently of the classical macroautophagy machinery (FIP200) but dependent on ATG16L1^16^. Thus, STING intracellular trafficking and signaling activities are tightly connected.

STING post-golgi trafficking to the endolysosomal compartment is essential for its degradation and signaling shutdown, but the proteins governing these processes and the signals triggering activated STING elimination require further investigation^17, 18^. Here we used a systems approach to identify genes that mediate STING trafficking. We show that oligomerization drives STING ubiquitination and that ubiquitinated STING recruits ESCRT in the endosomal compartment to achieve STING degradation and signaling shutdown. By focusing on genes mutated in human disease, we show that a pathogenic mutant of the ESCRT-I subunit UBAP1 blocks STING degradation and leads to accumulation of STING in the endolysosomal compartment at steady state, driving spontaneous activation of the sensor. Based on these findings we propose an updated model of STING trafficking and degradation.

## Results

### A time-resolved map of STING trafficking

To unbiasedly establish the hierarchy of molecular mechanisms governing STING trafficking and to identify proteins that interact with STING during this process, we used recently developed proximity ligation technologies that overcome the challenge of identifying native interactions by classical co-immunoprecipitation. We fused STING to the biotin ligase TurboID^19^ (Fig. 1a) for rapid (i.e. 10 minutes) ligation of biotin to proteins within a 10nm radius, followed by enrichment of labeled proteins via streptavidin pull-down. To verify that the construct was functional, we performed microscopy in 293T cells stably expressing STING-TurboID and confirmed STING translocation upon cGAMP stimulation and co-localization with streptavidin staining (Fig. S1a). Moreover, streptavidin pulldown following a cGAMP stimulation time-course showed time-dependent TBK1 labeling with signal increasing at 30 minutes, peaking at 2 hours and decreasing at 6 hours (Fig. 1b). Finally, the decrease in STING levels at the 6 hour time point indicated that this time course covers the full trafficking pathway from the ER to the site of STING degradation at the lysosome.

**Figure 1.**
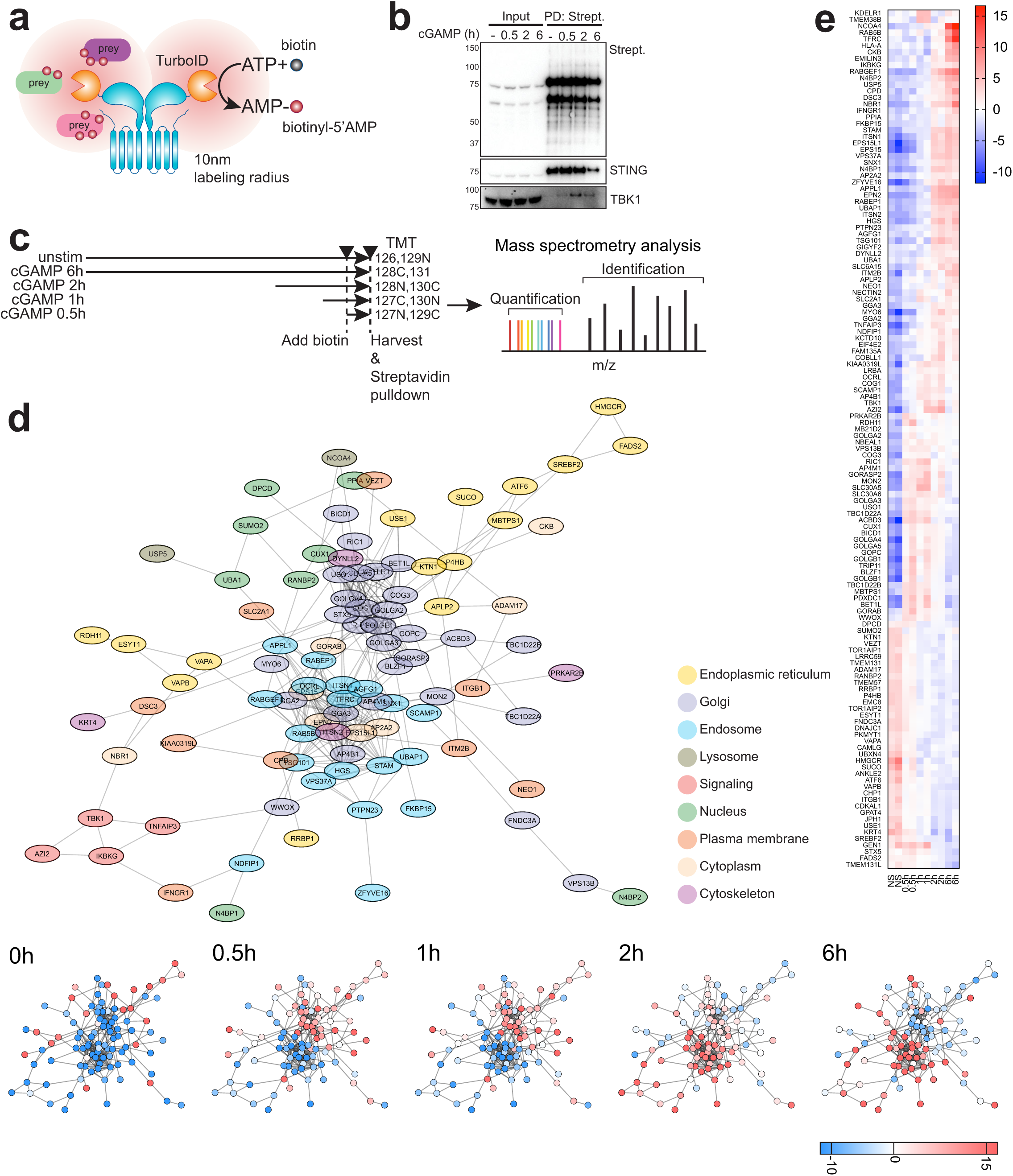
A time-resolved map of STING trafficking. **a)** Schematics of STING-TurboID fusion. STING was fused to TurboID at the CTT. Addition of biotin allows labeling of neighboring proteins in a 10nm radius. Labeled proteins can be then enriched after cell lysis with streptavidin pulldown. **b)** Immunoblot of Streptavidin-HRP (Strept.), STING and TBK1 in input and post streptavidin pull-down (PD: strept.) after 2µg/ml cGAMP stimulation (in perm buffer) for the indicated times in 293T STING-TurboID. **c)** Scheme of the time-course used for STING-TurboID proteomics. Reporter TMT labeling ions used for each condition are indicated. **d)** STRING generated network of filtered STING interactors after statistical analysis (top) and relative enrichment at the different time points (bottom). Colors in the top represent annotation of cellular compartments. Proteins were filtered on adj. p_value_<0.07 to include TBK1 (adj. p_value_=0.0611). **e)** Heat-map of the filtered proteins with enrichment at the different timepoints. n=2 per time-point.

To map protein-protein interactions, we next performed proximity labeling followed by quantitative mass spectrometry analysis. STING-TurboID expressing cells were stimulated with cGAMP and then were biotin-labeled for 30 minutes at different time points post-stimulation (Fig. 1c). Streptavidin enriched lysates were labeled with MS-differentiable tandem mass tags (TMT) and analyzed by liquid-chromatography tandem mass spectrometry (LC-MS/MS). Over 2000 proteins were identified in total across the different time-points. Fuzzy c-means clustering identified three protein clusters. We analyzed clusters and single timepoints to identify enriched Gene Ontology (GO) and Reactome terms (Fig S1b-d, S2a-d). Cluster 1 included proteins of the Golgi apparatus and vesicle transport, and showed an enrichment peak at 30 minutes and 1 hour with decreasing enrichment after the 2 hour time point (Fig. S1b, S2a-b). Cluster 2 contained endosomal proteins initially enriched at the 2 hour time point and slightly decreased at the 6 hour time point (Fig. S1c). At 2 hours, proteins were associated with “trans-Golgi network Vesicle Budding” and “Cargo recognition for clathrin mediated endocytosis”, indicative of STING exit from the Golgi through the TGN and trafficking to endosomes (Fig. S2c). In addition, at 2 hours, hits were associated with the endosome and ESCRT machinery and at 6 hours with the autolysosome and more weakly with ESCRT (Fig. S2c, S2d). Finally, cluster 3 included proteins of the ER that were enriched at time 0 (not stimulated) and decreased at all other time points (Fig. S1d).

Out of the ∼2000 proteins identified, statistical analysis led us to filter the dataset down to a network of 132 proteins that were differentially biotinylated between time-points. The inferred localization and trafficking pattern of this network of proteins was consistent with the analyses performed on the full data-set (Fig. 1d, 1e, S1b-d, S2), with ER proteins enriched at time 0, Golgi proteins at 30 minutes peaking at 1 hour, and endosome proteins peaking at 2 hours and still highly abundant at 6 hours. We identified a signaling hub comprising TBK1, AZI2, TNFAIP3 and IKBKG that was mostly enriched at two hours. Components of the ESCRT machinery were also abundant at the 2 hour time-point. NCOA4 and TFRC, involved in ferritin turnover through selective autophagy and highly enriched in lysosomes, were labeled at 6 hours, along with the endosomal marker RAB5B^20^. Finally we found two selective autophagy receptors, NBR1 enriched at 2 hours in the filtered dataset, and p62 enriched at 2 and 6 hours in the full dataset (Fig. 1e, S1e), indicative of a possible cross-talk between autophagy and STING trafficking to the endosome.

This dataset provides a time-resolved map of potential STING interactors and the basis to identify the hierarchy of regulators involved in STING trafficking and degradation.

### A genome-wide CRISPR screen identifies the HGS and VPS37A ESCRT subunits as required for STING degradation

To separately identify proteins required for STING trafficking and degradation, we performed a genome wide CRISPR screen based on activity of a STING protein reporter that undergoes degradation. We optimized a system to follow STING degradation with flow cytometry by fusing STING to the bright green fluorescent protein mNeonGreen (mNG) under control of a weak promoter. Stimulation of 293T cells expressing the reporter with the stable cGAMP analog 2’3’-cGAMP(pS)2 led to a decrease in STING-mNG signal (Fig. 2a), thus confirming functionality of the reporter cell line.

**Figure 2.**
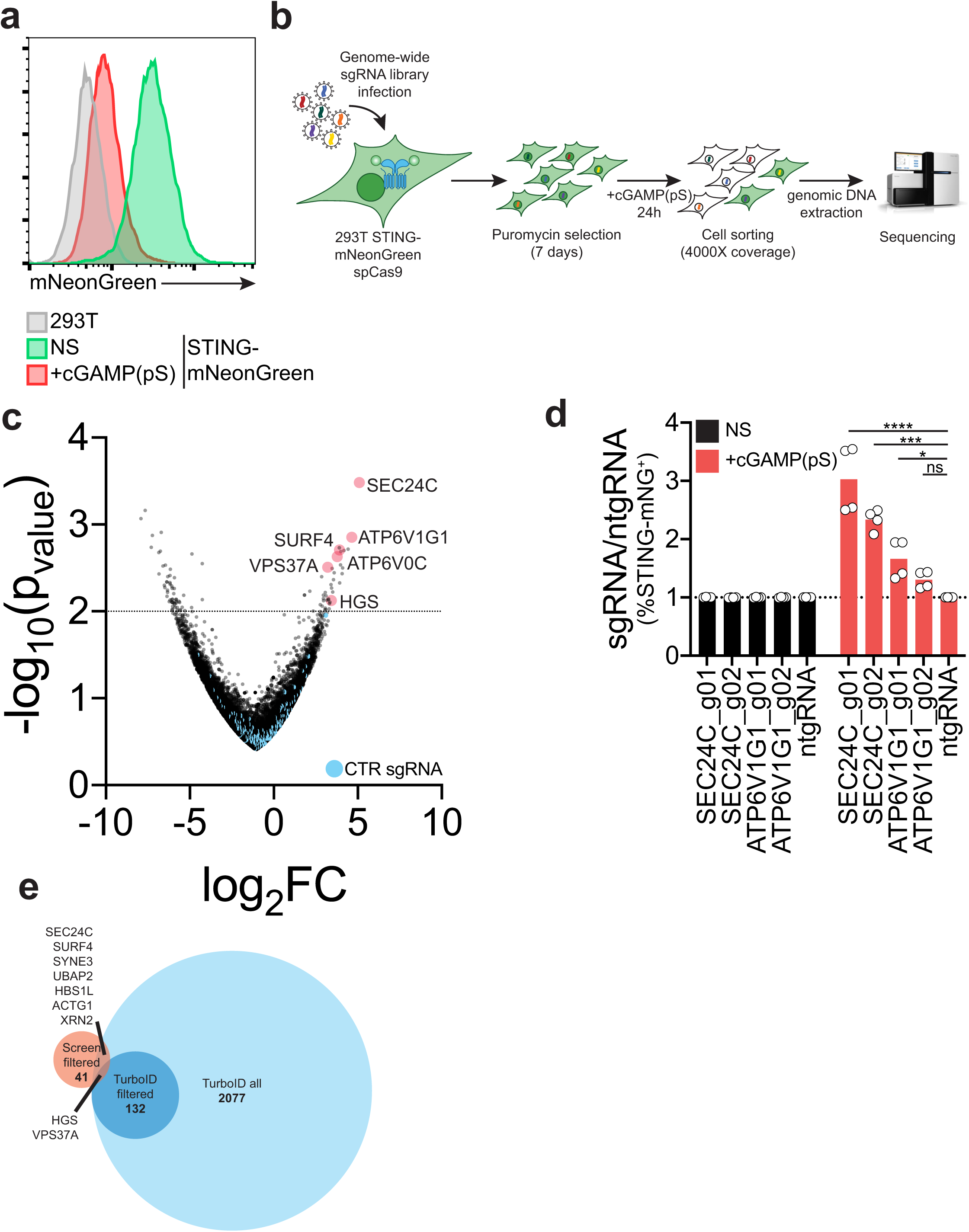
A genome-wide CRISPR screen identifies the HGS and VPS37A ESCRT subunits as required for STING degradation. **a)** mNeonGreen (mNG) signal intensity in 293T (gray) or a 293T cell line stably expressing a STING-mNeonGreen reporter non-stimulated (NS) (green) or stimulated (red) with 1µg/ml 2’3’-cGAMP(pS)2 (in medium) for 24 hours. One representative experiment of n=3 experiments. **b)** Strategy for genome-wide CRISPR screen to identify regulators of STING trafficking and degradation. **c)** Volcano-plot of log_2_ fold change (log_2_FC) vs -log_10_(p_value_) after sequencing and analysis of the genome-wide CRISPR screen as in b). Genes of interest are highlighted in red. Control guides are in blue. **d)** Percentage of STING-mNG positive 293T STING-mNG in cells stimulated or not with 1µg/ml 2’3’-cGAMP(pS)2 (in medium) for 24 hours. Showed is ratio %STING-mNG positive of each sgRNA over %STING-mNG positive cells of the control non-targeting sgRNA (ntgRNA). Two independent sgRNAs per gene. n=2 independent experiments with n=2 technical replicates per experiment. Each dot represents an individual replicate. One-way ANOVA with Dunnet’s multiple comparisons test. ****p<0.0001, ***p<0.001, *p<0.05, ns: not-significant. **e)** Intersection of all proteins identified by STING-TurboID proteomics (TurboID all) and filtered proteins (TurboID filtered) with hits from the genome-wide CRISPR screen with log_2_FC>0 and -log_10_(p_value_)>2.

To facilitate the CRISPR screen, we used a cell line stably expressing spCas9 and STING-mNG (Fig. 2b) that was transduced with the pooled genome-wide human-targeting sgRNA library Brunello^21^. One-week post selection for transduced cells, we stimulated the cells with cGAMP(pS)2, FACS-sorted mNG positive and negative cells (Fig S3a) and extracted genomic DNA to measure sgRNA abundance in the sorted populations by next-generation sequencing (Fig. 2b). When filtered, we found 41 positive regulators of degradation. The top scoring genes required for STING degradation were SEC24C and ATP6V1G1 (Fig. 2c). SEC24C was already described to regulate STING exit from the ER^8^ and ATP6V1G1 is a lysosomal V-ATPase subunit required for lysosomal acidification that was previously shown to be involved in STING degradation^17^. Individual sgRNA knock-outs for SEC24C and ATP6V1G1 recapitulated the results of our screen (Fig. 2d, S3b), validating our approach.

To determine which screen hits likely interact with STING, we intersected significant hits from the CRISPR screen with the full dataset of STING-TurboID-labeled proteins (Fig. 2e). In addition to SEC24C, we found SURF4, which was recently implicated in STING trafficking^22^, as well as SYNE3 which connects the cytoskeleton to the nucleus, the actin gene ACTG1, two genes involved in RNA processing, XRN2 and HBS1L, and the gene of unknown function UBAP2.

When intersected with the filtered 132 proteins from the TurboID dataset, we identified 2 genes, HGS and VPS37A, which are components of the ESCRT machinery (Fig. 2e). Overall, the intersection of our proteomics dataset with our genetic screen nominates candidate genes that potentially interact with STING and are required in regulation of its trafficking. We focused our attention on the two components of the ESCRT machinery identified, HGS and VPS37A.

### An ESCRT complex containing HGS and VPS37A regulates STING degradation and signaling shutdown

The ESCRT machinery is involved in many cellular processes that entail inverse membrane involution and formation of vesicles. Related to intracellular protein trafficking, ESCRT has been characterized to be required for intraluminal vesicles (ILVs) formation at late endosomes and more recently for resolution of particular forms of autophagy^23^. In addition to HGS (ESCRT-0) and VPS37A (ESCRT-I), the proteomics dataset identified other components of the ESCRT machinery: STAM (ESCRT-0), two subunits of the ESCRT-I heterotetrameric complex, TSG101 and UBAP1, and the Bro1 domain protein PTPN23 which has been shown to bridge ESCRT-I to ESCRT-III^24^ (Fig. 3a). Interestingly, UBAP1 has been shown to be preferentially assembled in an endosome-specific ESCRT complex containing VPS37A, but not its homologs VPS37B, C or D^25^.

**Figure 3.**
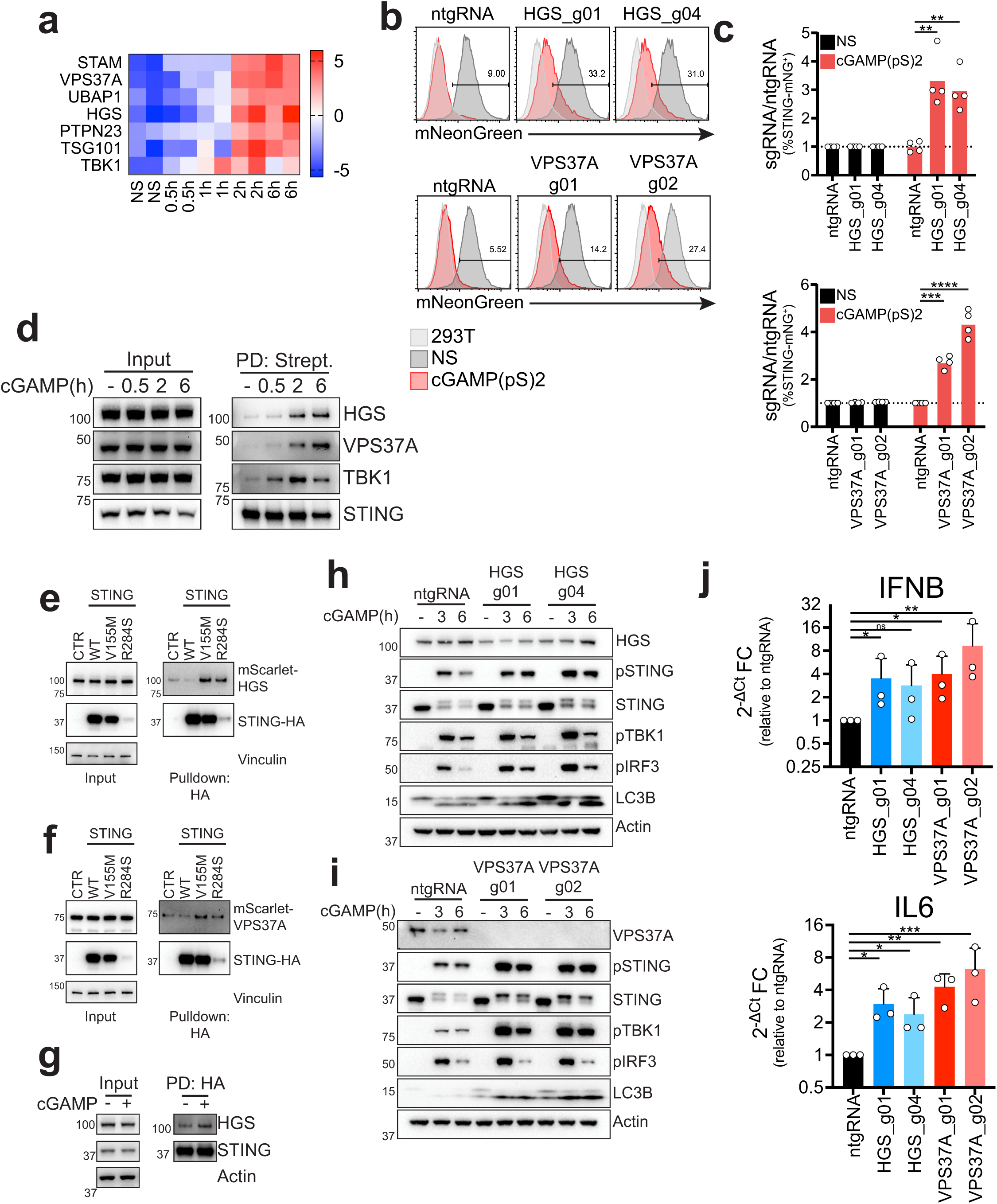
An ESCRT complex containing HGS and VPS37A regulates STING degradation and signaling shutdown. **a)** Heat-map of enriched proteins of the ESCRT machinery and TBK1 identified in the filtered STING-TurboID dataset. **b)** mNeonGreen levels in 293T STING-mNeonGreen cell lines KO for the indicated genes before (dark gray – NS) or after (red) stimulation with 1µg/ml 2’3’-cGAMP(pS)2 (in medium) for 24 hours. Line represents gating strategy and numbers represent %STING-mNG positive cells post stimulation. 293T (light gray) are shown as a reference for mNG negative cells. One representative plot of n=2 independent experiments with n=2 technical replicates per experiment. **c)** Percentage of STING-mNeonGreen (mNG) positive cells in cells stimulated or not (NS) with 1µg/ml 2’3’-cGAMP(pS)2 (in medium) for 24 hours. Shown is ratio %STING-mNG positive of each sgRNA over %STING-mNG positive cells of the control non-targeting sgRNA (ntgRNA). Two independent sgRNAs per gene. n=2 independent experiments with n=2 technical replicates per experiment. Each dot represents an individual replicate. One-way ANOVA with Dunnet’s multiple comparison test. ****p<0.0001, ***p<0.001, **p<0.01. **d)** Immunoblot of the indicated proteins in 293T STING-TurboID stimulated with 2µg/ml cGAMP (in perm buffer) for the indicated times, in the input and after streptavidin pulldown (PD: Strept.). One representative blot of n=3 independent experiments. **e)** Immunoblot of the indicated proteins in the input and post HA co-immunoprecipitation in 293T cells stably transduced with either an empty vector (CTR) or the indicated STING mutants, after transfection with an mScarlet-HGS expressing vector. One representative blot of n=3 independent experiments. **f)** Same as in e) but cells were transfected with mScarlet-VPS37A. **g)** Immunoblot of the indicated proteins in the input and post HA co-immunoprecipitation in 293T cells stably transduced STING-HA, after transfection with a mScarlet-HGS expressing vector stimulated or not with 2µg/ml cGAMP (in perm buffer) for 3 hours. One representative experiment of n=2 independent experiments. **h)** Immunoblot of the indicated proteins in BJ1 fibroblasts KO for HGS two independent guides per gene or transduced with a non-targeting sgRNA (ntgRNA). Cells were stimulated with 0.5µg/ml cGAMP (in perm buffer) for the indicated times. One representative blot of n=3 independent experiments. **i)** Same as in g for VPS37A. **j)** qPCR for IFNβ (top) and IL6 (bottom) in BJ1 fibroblasts KO for HGS (blue) or VPS37A (red) stimulated with 0.5µg/ml cGAMP (in perm buffer) for 8 hours. Shown is 2^-ΔCt^ Fold Change calculated as ratio 2^-ΔCt^ sgRNA/2^-ΔCt^ ntgRNA for cells stimulated with cGAMP. One-way ANOVA on log-transformed data with Dunnet multiple comparison test. *p<0.05, **p<0.01, ***p<0.001, ns=not significant.

We focused our attention on HGS and VPS37A because they were identified at the intersection of our proteomics and genetics data. When knocked out with two independent sgRNAs per gene, KO STING-mNG reporter cells showed a reduction in STING degradation, confirming our screen results (Fig. 3b-c, S4a-b). STING-TurboID mediated labeling of HGS and VPS37A started at 2 hours post cGAMP stimulation and was stable up to 6 hours, confirming our mass-spectrometry data (Fig. 3d). Since proximity ligation identifies interacting and non-interacting proteins in the proximity of the bait, we also performed co-immunoprecipitations (co-IP) with STING. We generated 293Ts stably expressing a control vector, HA-tagged wild-type (WT) STING, or the constitutive active mutants STING V155M and STING R284S found in SAVI patients^26–28^. HGS and VPS37A co-immunoprecipitated with the STING constitutive active mutants V155M and R284S, but not with WT STING (Fig. 3e, 3f). In addition, stimulation of WT STING expressing cells led to increased pulldown of HGS (Fig. 3g). This confirmed a physical interaction between activated STING and the ESCRT subunits HGS and VPS37A.

Since STING functions are dependent on its intracellular trafficking and degradation, we asked if HGS and VPS37A KO would impact STING signaling. We generated CRISPR KO human primary fibroblasts (BJ1) with two independent sgRNAs per gene. KO of HGS or VPS37A increased STING signaling, phosphorylation of TBK1 and IRF3, (Fig. 3h-i), decreased STING degradation and induced increased transcription of IFNβ and IL6 (Fig. 3j, S4c) without altering total TBK1 and IRF3 levels (Fig. S4d-e). Impaired STING degradation and failure to shut-down STING signaling leads to cell death^29^. The U937 monocytic cell line has been shown to be susceptible to STING dependent cell death^30^. KO of HGS and VPS37A in U937 cells led to increased STING signaling in response to cGAMP (Fig. S4f-g) resulting in increased cell death including higher levels of Annexin V staining 24h post-treatment (Fig. S4h-k).

Finally, since resolution of membrane involution by ESCRT is dependent on VPS4A/B, we tested if STING degradation was dependent on these proteins. We overexpressed the VPS4A E228Q dominant negative (DN) mutant fused to mScarlet in a STING-HA reporter cell line. When stimulated with cGAMP, there were comparable levels of STING degradation between non transfected cells and cells transfected with a control plasmid (Fig. S4l-n), while cells expressing the VPS4A DN showed a marked reduction in STING degradation (Fig. S4l-n), consistent with an ESCRT requirement for this process.

Taken together, these data suggest that ESCRT is required for both STING degradation and signaling shut-down.

### ESCRT links STING degradation and autophagy resolution at the endosome

Interestingly, HGS and VPS37A KO in fibroblasts not only blocked STING degradation but also increased lipidated LC3B levels, an autophagy marker that accumulates in cells with defective resolution of autophagy^31^ (Fig. 3h-i). This finding led us to ask if the endosome and the autophagy degradative pathways, previously proposed to be distinct and to proceed respectively from the TGN or from the ERGIC^8^, were instead part of one coordinated shutdown mechanism. The ESCRT-I component VPS37A, which acts downstream of the ESCRT-0 subunit HGS, has recently been shown to be required for phagophore closure in macroautophagy^32^. Consistent with these findings, VPS37A KO fibroblasts showed a low basal level of accumulation of lipidated LC3B in absence of cGAMP (Fig. 3i).

To visualize intracellular trafficking of STING, we stained cells for the ESCRT-0 subunit HGS (which acts upstream of VPS37A), the late endosomal/lysosomal marker CD63 and the autophagy markers p62 (identified in our proteomics), or LC3B. Upon cGAMP stimulation, HGS formed distinct foci that co-localized with STING and were in close proximity to CD63, indicating that STING and ESCRT co-localize at the late endosome/lysosome (Fig. 4a, Fig. S5a). We then looked at co-localization of STING and the autophagy receptor p62, which was identified in our proteomics. Similar to HGS, p62 formed distinct foci in the cell upon cGAMP stimulation (Fig. 4b, S5b). p62 foci colocalized with HGS and STING, suggesting that ESCRT could be involved in resolution of STING induced autophagy. Interestingly, when we could identify clear circular p62 structures, we noticed that HGS and STING formed a bright signal at one distinct focus on these structures (Fig. 4b). Similar circular structures with a bright spot of STING and HGS colocalization were identified when we stained for LC3B (Fig. 4c, S5c).

**Figure 4.**
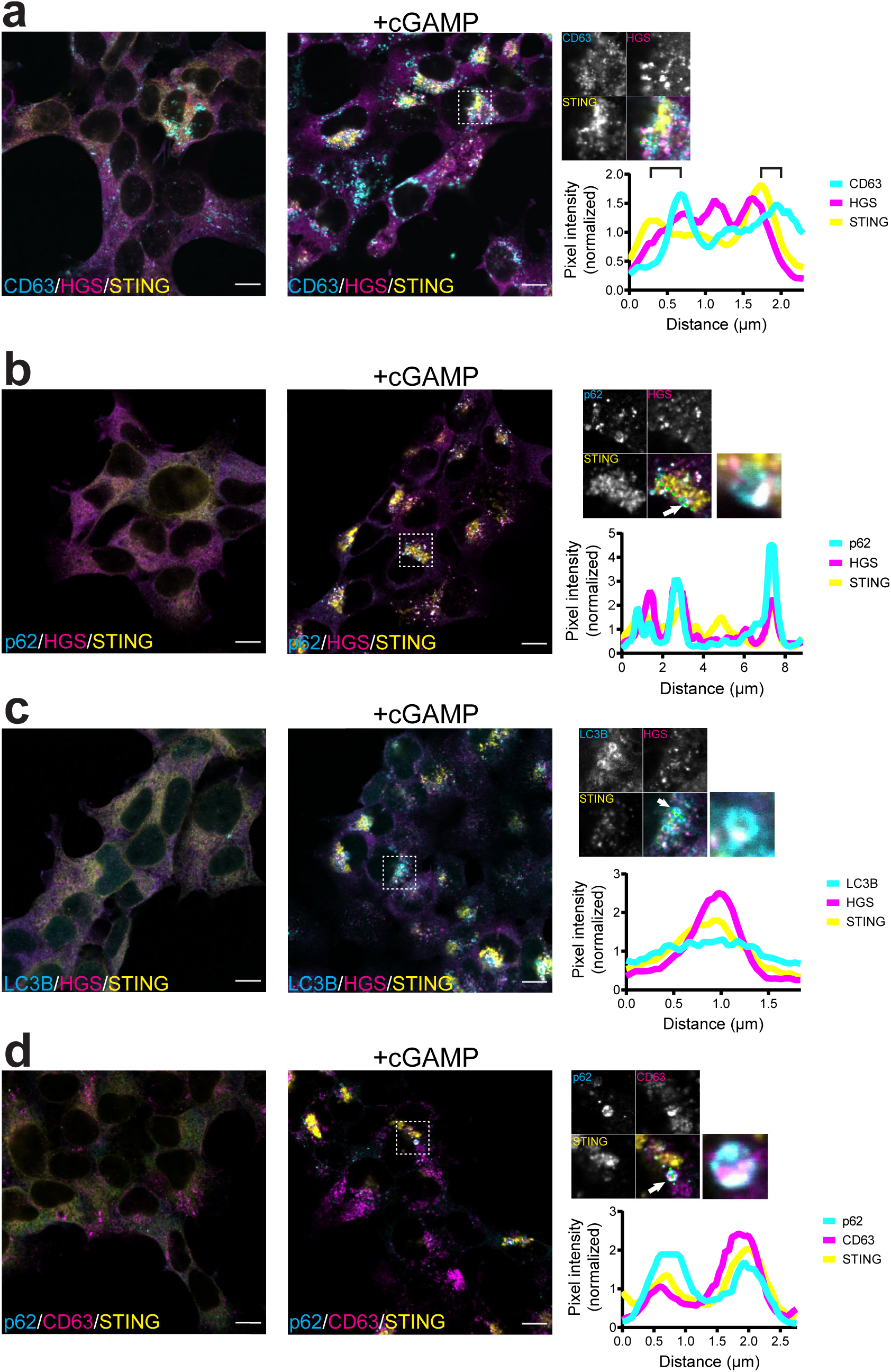
ESCRT links STING degradation and autophagy resolution at the endosome. Immunofluorescence of **a)** CD63 (cyan), HGS (magenta) and STING (yellow), **b)** p62 (cyan), HGS (magenta) and STING (yellow), **c)** LC3B (cyan), HGS (magenta) and STING (yellow), **d)** p62 (cyan), CD63 (magenta) and STING (yellow) in 293T stably expressing STING-HA non stimulated (left panels) or after stimulation with 2µg/ml cGAMP (in perm buffer) for 2 hours (right panels). Dashed box represents the cropped region shown in the enlarged panels. Green dashed line represents the line used to plot normalized pixel intensity for each protein. Brackets in the profile in panel a represent proximity of STING/HGS peaks to CD63 peaks. Panels b, c and d show also an enlargement of the structures indicated by the white arrow. One representative field of n≥3 fields in n=3 independent experiments. Scale bar is 10µm. Corresponding single color images are in Fig. S5.

The co-localization of STING and ESCRT in proximity to the late endosomal/lysosomal marker CD63 and the autophagy markers p62 and LC3B prompted us to ask if resolution of STING-dependent autophagy could happen at the late endosome/lysosome. LC3B (encoded by *ATG8*) has been shown to be conjugated to single membranes upon STING activation in a process called conjugation of ATG8 in the endolysosomal compartment (CASM)^16, 33^. If STING containing vesicles in the endosomal compartment are the target of LC3B conjugation and are degraded via ESCRT mediated fusion at the late endosome, the autophagy marker p62 and CD63 would colocalize and accumulation of LC3B in ESCRT KO fibroblasts would derive from failure of degradation. We stained STING expressing cells for p62 and CD63 (Fig 4d, S5d) and indeed observed co-localization of the two markers when cells were treated with cGAMP.

Taken together, these data suggest that vesicles containing activated STING are LC3B lipidated and marked with p62. ESCRT-dependent fusion of these vesicles with the late endosome/lysosome leads to STING degradation and consequent reduction in lipidated LC3B.

### STING ubiquitination creates a platform at the endosome for STING degradation

ESCRT-0 and ESCRT-I are known to associate with ubiquitinated cargo at the endosome, and HGS and VPS37A both contain a ubiquitin binding motif^23^. We asked if STING was ubiquitinated upon cGAMP stimulation and if ubiquitination could drive its association with ESCRT. To block ubiquitination of STING, we used the Ubiquitin Activating enzyme 1 (UBA1) inhibitor MLN7243^34^. MLN7243 blocked STING ubiquitination upon cGAMP stimulation (Fig. S6a). When we stimulated our STING-mNG reporter cell line with cGAMP, there was a complete block of degradation of STING in presence of the inhibitor (Fig. 5a-b). Ubiquitination can drive protein degradation through the proteasome or through intracellular protein trafficking to the late endosome. To discriminate between these two pathways, we treated STING-mNG expressing cells with two proteasome inhibitors, Bortezomib and MG-132, the UBA1 inhibitor MLN7243 and the lysosomal V-ATPase inhibitor Bafilomycin A1. Both proteasome inhibitors only partially impacted STING degradation, consistent with previous reports^17, 35^, while this process was completely blocked by both MLN7243 and Bafilomycin A1, suggesting that sorting to the late endosome and acidification of the lysosome are central to STING degradation (Fig S6b-c). Therefore, we hypothesized that MLN7243 blocked STING degradation by inhibiting its ubiquitination, consequently preventing its interaction with HGS and VPS37A. To test this hypothesis, we treated STING-TurboID expressing cells with MLN7243. Treatment with the drug reduced the biotin labeling of HGS and VPS37A, while labeling of TBK1 was unaffected (Fig. 5c). These results suggest that STING ubiquitination upon activation drives its association with ESCRT and regulates its degradation. We then asked if the UBA1 inhibitor MLN7243 would recapitulate HGS and VPS37A KO effects on STING signaling, degradation and autophagy. Indeed, treatment with MLN7243 abrogated STING degradation in fibroblasts, increased phospho-STING signaling, induced accumulation of lipidated LC3B (Fig. 5d) in addition to increased IFNβ and IL6 transcription (Fig. 5e).

**Figure 5.**
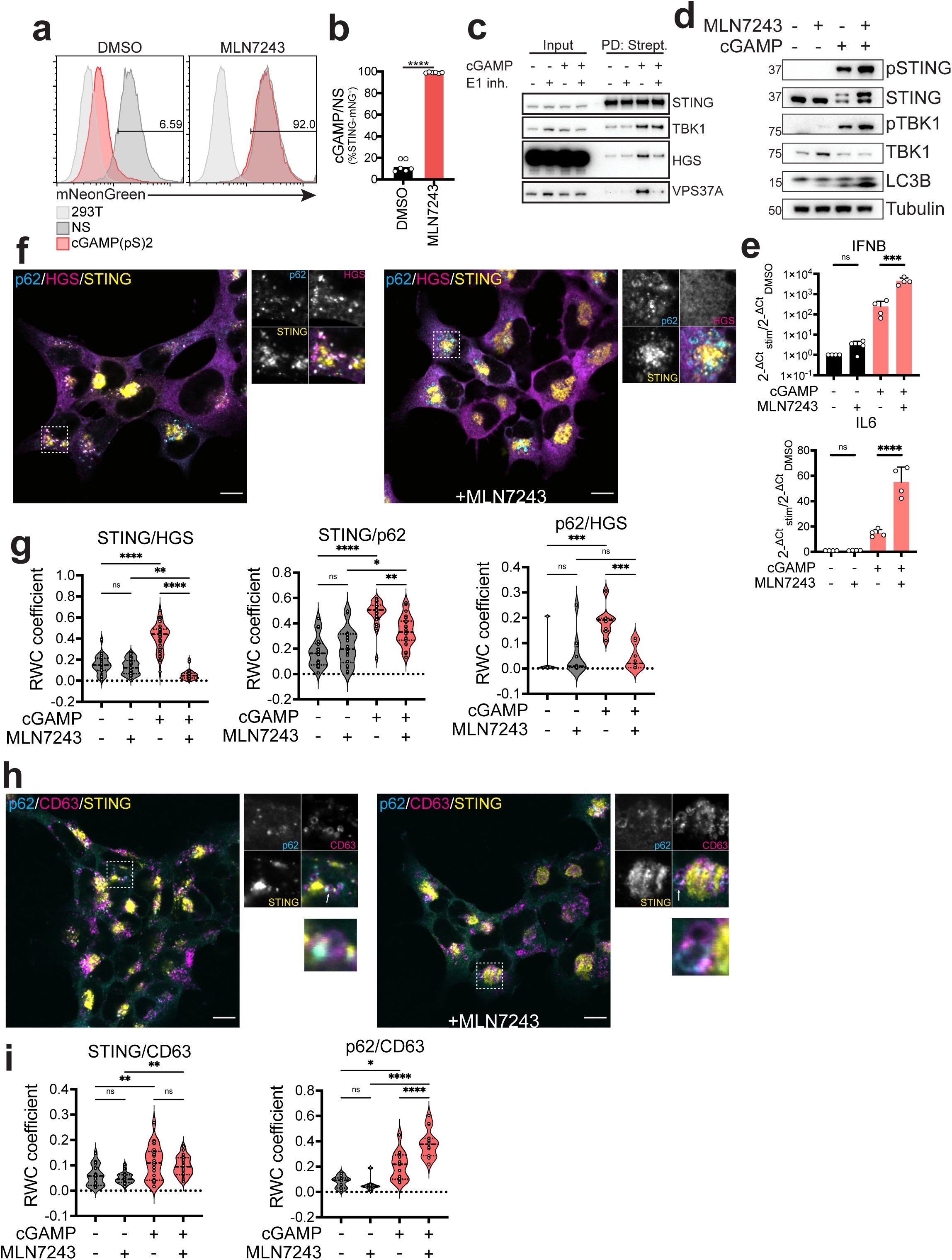
STING ubiquitination creates a platform at the endosome for STING degradation. **a)** mNeonGreen levels in the 293T STING-mNeonGreen reporter cell line before (dark gray – NS) or after (red) stimulation with 4µg/ml 2’3’-cGAMP(pS)2 (in medium) for 8 hours in presence or in absence of 0.5µM of the UBA1 inhibitor MLN7243. Line represents gating strategy and numbers represent %STING-mNeonGreen positive cells post stimulation (red). 293T (light gray) are shown as a reference for mNeonGreen negative cells. One representative plot of n=3 independent experiments with n=2 technical replicates per experiment. **b)** Percentage of STING-mNeonGreen (mNG) positive cells in cells stimulated or not as in a). Shown is ratio %STING-mNG positive post simulation over %STING-mNG positive cells of non-stimulated cells. n=3 independent experiments with n=2 technical replicates per experiment. Each dot represents an individual replicate. Paired t-test. ****p<0.0001 **c)** Immunoblot of the indicated proteins in the input or after streptavidin pull-down in 293T STING-TurboID cells stimulated or not with 2µg/ml cGAMP (in perm buffer) for 3 hours in presence or in absence of 0.5µM MLN7243. One representative blot of n=3 independent experiments. **d)** Immunoblot of the indicated proteins in BJ1 fibroblasts stimulated or not with 0.5µg/ml cGAMP (in perm buffer) for 2 hours in presence or in absence of 0.5µM MLN7243. One representative blot of n=3 independent experiments. **e)** qPCR for IFNβ (top) and IL6 (bottom) in BJ1 fibroblasts stimulated with 0.5µg/ml cGAMP (in perm buffer) for 8 hours with or without 0.5µM MLN7243. Shown is ratio 2^-ΔCt^ of each condition/2^-ΔCt^ of unstim treated with DMSO (cGAMP-/MLN7243-). n=2 independent experiments with n=2 technical replicates. One-way ANOVA on log-transformed data with Dunnet multiple comparison test. ***p<0.001, ****p<0.0001 ns=not significant. **f)** Immunofluorescence of p62 (cyan), HGS (magenta) and STING (yellow) in absence (left) or in presence (right) of MLN7243 in 293T stably expressing STING-HA after stimulation with 2µg/ml cGAMP (in perm buffer) for 2 hours. Dashed boxes represent the cropped regions shown in the right panels. One representative field of n≥5 fields in n=2 independent experiments. Scale bar is 10µm. Control non-stimulated cells are in Fig. S7. **g)** Rank Weighted Colocalization (RWC) coefficient for HGS colocalization in STING foci, STING colocalization in p62 foci and p62 colocalization in HGS foci in cells stimulated as in f. Each dot represents colocalization calculated in a field. All images in which STING was co-stained with a marker in any combination were aggregated. n=2 independent experiments with n≥5 fields. One way ANOVA with post-hoc Tukey test. *p<0.05, **p<0.01, ***p<0.001, ****p<0.0001, ns=not significant. **h)** Immunofluorescence of p62 (cyan), CD63 (magenta) and STING (yellow) in cells stimulated as in f). **i)** RWC of STING in CD63 foci and p62 in CD63 foci. Each dot represents colocalization calculated in a field. All images in which STING was co-stained with a marker in any combination were aggregated. n=2 independent experiments with n≥5 fields. One way ANOVA with post-hoc Tukey test. *p<0.05, **p<0.01, ***p<0.001, ****p<0.0001, ns=not significant.

We wanted to identify the impact of UBA1 inhibition on STING subcellular localization. When treated with MLN7243, STING showed a different intracellular distribution pattern lacking dispersion in perinuclear vesicles (Fig. 5f, 5h, S6d, S6f, S7a-d) and HGS foci co-localizing with STING were completely lost (Fig. 5f-g, S7a). HGS also failed to co-localize with p62 in presence of MLN7243 (Fig. 5f-g, S7a). The same was true for co-localization with CD63 and HGS (S6d-e, S7c). Consistent with treatment in fibroblasts (Fig. 5d), MLN7243 also led to an increase of STING colocalization with LC3B, while decreasing its colocalization with HGS (Fig. S6f-g). Moreover, we noticed that the distribution of p62 changed from one spot co-localizing with STING at one edge of CD63^+^ vesicles to an accumulation of p62^+^ ring-like structures, possibly indicative of LC3B lipidated STING vesicles failing to fuse with the late endosome (Fig. 5f, 5h, S7a-b), reflected in increased colocalization between p62 and CD63 (Fig. 5i). In presence of MLN7243, we identified p62^+^ vesicles in the process of fusing with CD63^+^ vesicles (Fig. 5h) suggesting that STING-induced ESCRT-dependent resolution of autophagy through fusion at the endolysosomal compartment was blocked.

Taken altogether, we hypothesize that, analogous to phosphorylation that creates a platform at the TGN for type I IFN signaling, ubiquitinated STING traffics to and decorates the endosome creating an organizing platform that coordinates STING degradation and autophagy shutdown through ESCRT.

Mutations in UBA1 have been recently identified in patients, leading to an autoinflammatory disease named Vacuoles E1 enzyme X-linked Autoinflammatory Somatic (VEXAS) syndrome^36^. VEXAS syndrome patients express a *de novo* hypomorphic UBA1 isoform in the myeloid compartment. Since these patients show type I IFN and proinflammatory cytokine production at steady state, we wondered if UBA1 inhibition in CD14^+^ monocytes could lead to an increase in STING signaling. After cGAMP stimulation, CD14^+^ monocytes from healthy donors showed a strong increase in phospho-STING levels when UBA1 was inhibited by MLN7243 treatment (Fig. S6h). Human monocytes have been shown to activate NLRP3 dependent pyroptosis in response to STING ligands due to lysosomal membrane permeabilization leading to IL1β release^37^. We hypothesized that the exacerbated STING signaling induced by MLN7243 in these cells could lead to exacerbated cell death. Indeed, MLN7243 synergized with cGAMP to induce cell death (Fig. S6i-j). Since VEXAS is an autoinflammatory disease with late onset (median age at onset – 64), these results might suggest that the inflammation in these patients could be driven by increased STING signaling due to defective degradation triggered by an increased presence of cytosolic DNA associated with cellular senescence that would lead to increase in IL1β release^38–40^.

### A targeted CRISPR screen identifies UBE2N as a regulator of STING degradation

To specifically identify ubiquitin related genes involved in STING degradation, we performed a targeted CRISPR screen in both the STING-mNG and STING-HA reporter cell lines. The library contained guides targeting 669 E3 and adaptors, 40 E2, 7 E1, 28 Autophagy core proteins and 10 positive controls from the genome-wide CRISPR screen. While the STING-mNeonGreen performed better than the STING-HA screen, both recovered positive controls as required for STING degradation (SEC24C, ATP6V1G1, ATP6V0C, HGS, VPS37A, UBAP1) and STING as the top depleted gene (Fig. S8a). To increase our confidence in identifying relevant genes, we correlated both screens based on average fold change (Fig. 6a). Positive controls correlated strongly in both screens. Interestingly, we were able to clearly identify genes involved in autophagy as required for STING degradation, with ATG9A, ATG12, ATG5 and ATG16L1 highly enriched in both screens, suggesting that autophagy plays a role in STING degradation contrary to previous reports^8^. Indeed, KO of ATG16L1 and ATG5 in STING-mNG cells agreed with the screens results (Fig. S8b-d). Additionally, KO of ATG16L1 and ATG5 in BJ1 fibroblasts led to complete loss of LC3B lipidation, in accordance with previous findings, but also to a reduction of STING degradation and increase in STING signaling, contrary to what previously described^8, 16^ (Fig. S8e-f). When we looked at ubiquitin related genes, we focused our attention on the E2 conjugating enzyme UBE2N. UBE2N was present in our TurboID dataset and showed a peak enrichment at 1h post cGAMP stimulation that rapidly dropped at the 2 hour time-point, preceding association of STING with ESCRT (Fig. 6b). We confirmed STING-TurboID labeling of UBE2N via western blot (Fig. S8g). Based on our proteomics (Fig. 1d, S2), these results suggest that UBE2N is in proximity of STING at the Golgi (1h time-point) and is rapidly dissociating from STING post-Golgi.

**Figure 6.**
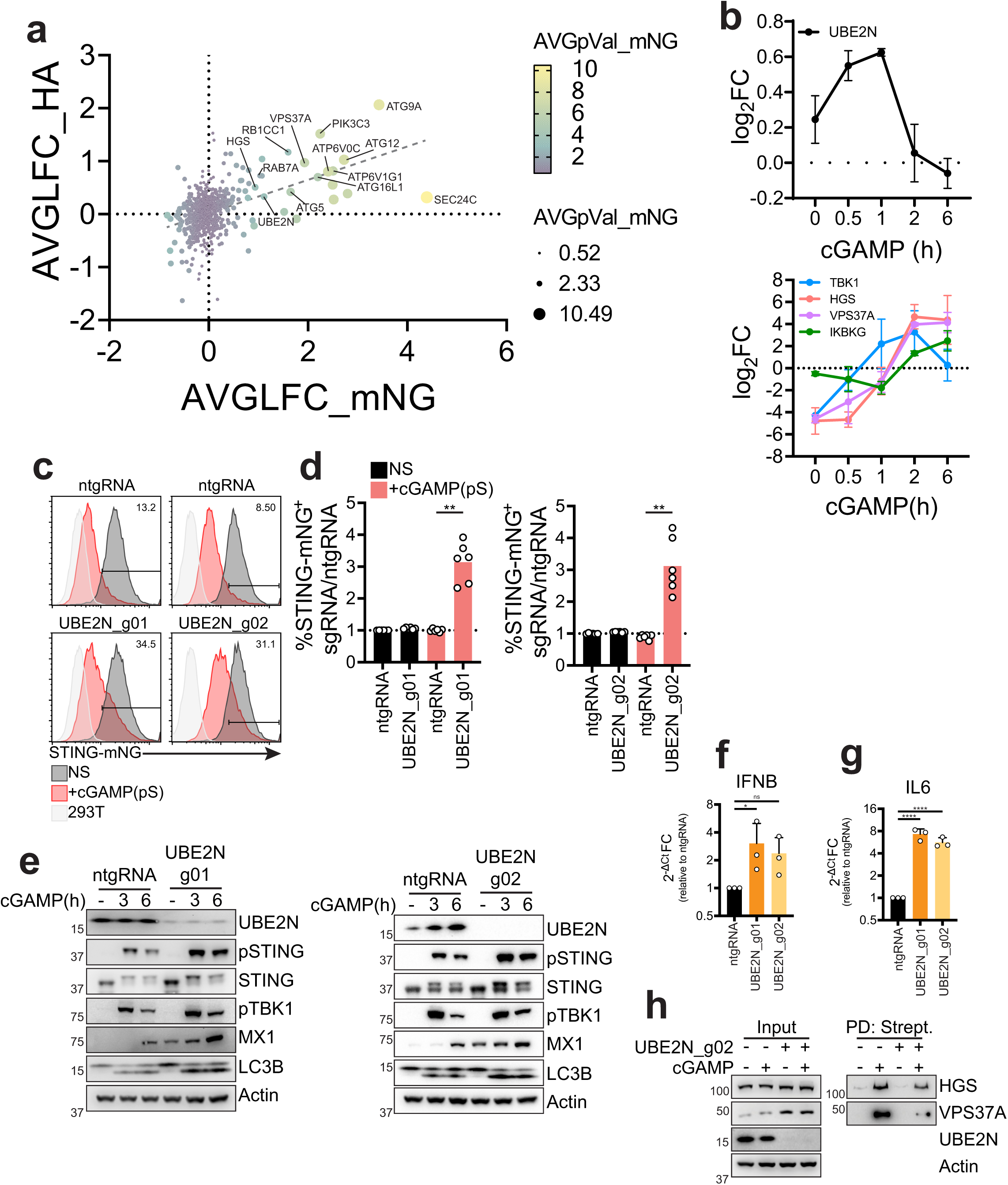
A targeted CRISPR screen identifies UBE2N as a regulator of STING degradation. **a)** Correlation plot of Average Log2 Fold-Change (AVGLFC) for the CRISPR screen in STING-mNG or STING-HA cell lines. Color and size represent Average p-value (AVGpVal) for the screen in mNeonGreen cells. **b)** log2 fold-change (FC) enrichment of the indicated proteins in TurboID proteomics described in Fig. 1. **c)** Percentage of STING-mNG positive cells in cells stimulated or not with 4µg/ml 2’3’-cGAMP(pS)2 (in medium) for 6 hours transduced with the indicated sgRNAs. One representative plot of n=3 experiments with n=2 technical replicates per experiment. **d)** Ratio %STING-mNG positive of each sgRNA over %STING-mNG positive cells of the control non-targeting sgRNA (ntgRNA) in cells stimulated as in c). n=3 independent experiments with n=2 technical replicates per experiment. Each dot represents an individual replicate. One-way ANOVA with Dunnet multiple comparisons test. **p<0.01. **e)** Immunoblot of the indicated proteins in BJ1 fibroblasts stimulated with 0.5µg/ml cGAMP (in perm buffer) for the indicated times. **f)** qPCR for IFNβ in BJ1 fibroblasts stimulated with 0.5µg/ml cGAMP (in perm buffer) for 8 hours. Shown is 2^-ΔCt^ Fold Change (FC) calculated as ratio 2^-ΔCt^ sgRNA/2^-ΔCt^ ntgRNA for cells stimulated with cGAMP. n=3 independent experiments. One-way ANOVA on log-transformed data with Dunnet multiple comparison test. *p<0.05, **p<0.01, ***p<0.001, ns=not significant. **g)** Same as in f) for IL6. **h)** Immunoblot of the indicated proteins in 293T STING-TurboID transduced with a control guide or knockout for UBE2N stimulated with 2µg/ml cGAMP (in perm buffer) for 3h.

UBE2N is responsible for K63 polyubiquitination of proteins that prompts them for degradation through the endolysosomal compartment and has been shown to play a regulatory role in multiple innate immune sensing pathways^41^. Consistent with a possible role for UBE2N in STING biology, STING has been shown in multiple studies to be K63 polyubiquitinated upon activation^35, 42, 43^. Consistent with these findings, NEMO (IKBKG), which interacts with K63 ubiquitin^44^, was found highly enriched in our TurboID dataset, following STING labeling of UBE2N (Fig. 6b). To validate the results of our screens, we knocked out UBE2N with two individual sgRNAs in 293T STING-mNG and stimulated the cells with cGAMP. UBE2N KO cells showed reduced STING degradation (Fig. 6c-d). We then knocked out UBE2N in fibroblasts to study the impact on STING signaling. Consistent with ESCRT KO, UBE2N KO led to decreased STING degradation, increased STING signaling, including increased IFNβ and IL6 transcription, and accumulation of lipidated LC3B (Fig. 6e-g, S8h-i). To test if the UBE2N KO effect on STING degradation and signaling was due to impaired STING association with ESCRT, we used the STING TurboID cell line in which we knocked-out UBE2N. KO of UBE2N led to a modest reduction of association with HGS upon cGAMP stimulation, while leading to a striking reduction of association with VPS37A (Fig. 6h). These results suggest that UBE2N activity regulates STING degradation through its association with ESCRT, impacting VPS37A recruitment to STING.

### STING oligomerization drives its ubiquitination and multiple lysines regulate STING degradation

Finally, we wanted to identify the trigger for STING ubiquitination. STING has been shown to oligomerize upon cGAMP binding^13, 45^. Protein aggregates have been shown to be cleared by association with the selective autophagy receptors p62 and NBR1^46^. In addition to p62, in accordance with our proteomics (Fig. 1e), STING colocalized with NBR1 and HGS upon activation (Fig. S9a). Association of p62 and NBR1 with cargo is ubiquitin dependent^47^ and treatment with MLN7243 leads to a reduction in STING colocalization with p62 (Fig. 5g). We then reasoned that STING oligomerization could drive its ubiquitination, consequently leading to its degradation. To block STING oligomerization without impacting STING exit from the ER, we used the STING palmitoylation inhibitor H-151^48^. Treatment of 293T STING-mNG cells with the inhibitor led to a marginal but significant decrease in STING degradation (Fig. 7a-7b). This effect was independent of TBK1 recruitment, since STING L374A^49^ did not show a difference in degradation compared to WT (Fig. S9b-c). To test if H-151 blocked STING ubiquitination, we used THP-1 expressing HA-ubiquitin and performed pulldowns. Indeed, treatment with H-151 led to reduced STING activation and ubiquitination reflected in a reduction in STING degradation to a level comparable to the effect seen in 293T STING-mNG (Fig. 7c).

**Figure 7.**
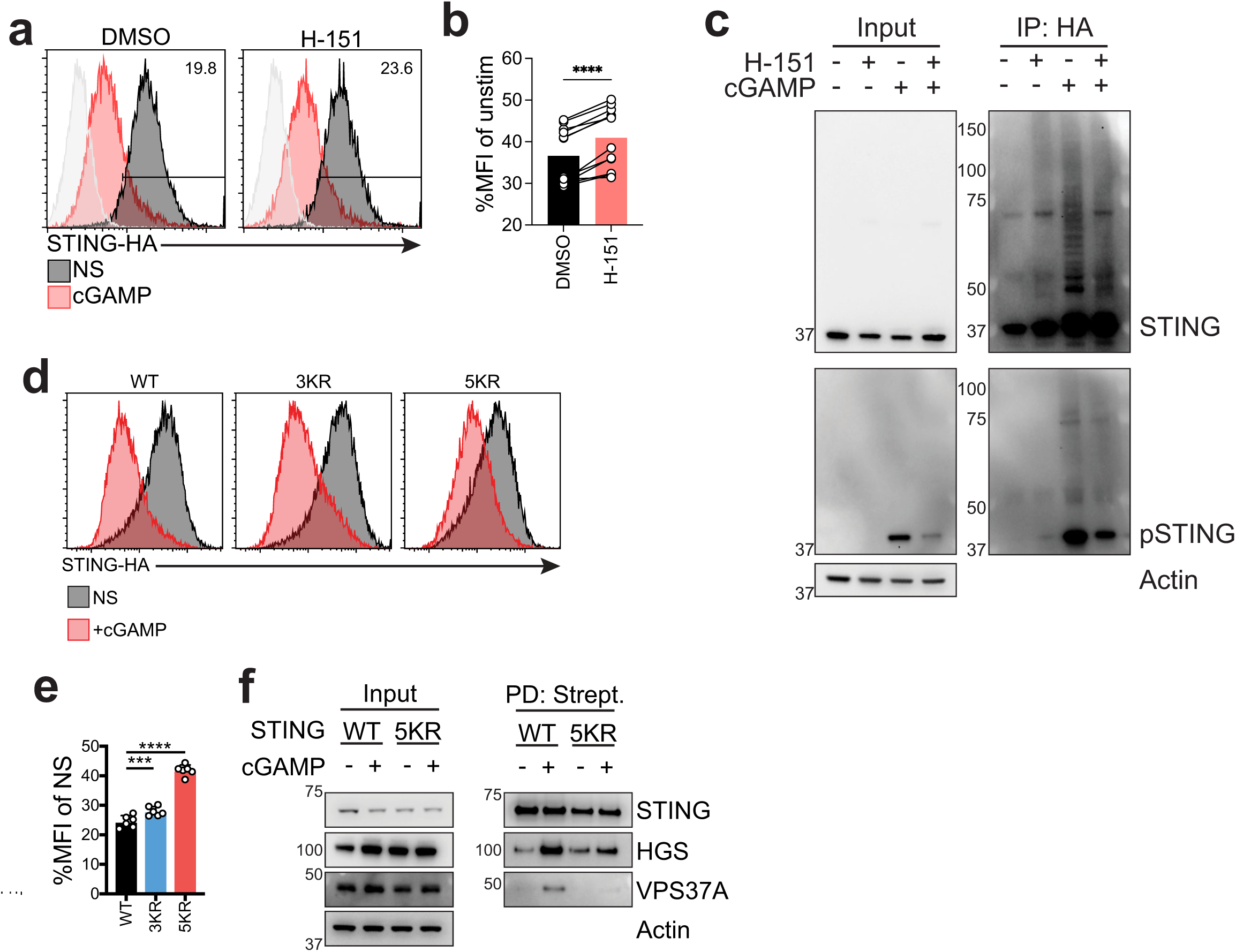
STING oligomerization drives its ubiquitination and multiple lysines regulate STING degradation. **a)** mNeonGreen levels in the 293T STING-mNeonGreen reporter cell line before (dark gray – NS) or after (red) stimulation with 4µg/ml 2’3’-cGAMP(pS)2 (in medium) for 8 hours in presence or in absence of 1µM H-151. Line represents gating strategy and numbers represent %STING-mNeonGreen positive cells post stimulation. 293T (light gray) are shown as a reference for mNeonGreen negative cells. One representative plot of n=4 independent experiments with n=3 technical replicates per experiment. **b)** Ratio MFI of experiment same as in a) shown as percentage of unstimulated. n=4 independent experiments with n=3 technical replicates per experiment. Each dot represents a replicate. Paired t-test. ***p<0.001. **c)** Immunoblot of the indicated proteins in THP-1 stably expressing HA-ubiquitin stimulated with 10µg/ml cGAMP (in medium with digitonin) for 4 hours. One experiment representative of n=3 independent experiments. **d)** HA intracellular staining in 293T stably expressing HA labeled STING WT, STING 3KR (K20R/K150R/K236R) or STING 5KR (K20R/K150R/K236R/K338R/K370R) mutants stimulated (red) or not (dark grey) with 2µg/ml cGAMP (in perm buffer) for 6 hours. One representative plot of n=3 independent experiments with n=2 technical replicates per experiment. **e)** MFI of cells as in d) shown as %MFI of cGAMP stimulated over non-stimulated (NS) for each mutant. n=3 independent experiments with n=2 technical replicates per experiment. Each dot represents an individual replicate. One-way ANOVA with Dunnet multiple comparisons. ****p<0.0001, ***p<0.001 **f)** Immunoblot of the indicated proteins in input and streptavidin pulldown from 293T cells expressing STING WT or STING 5KR fused to TurboID stimulated with 2µg/ml cGAMP (in perm buffer) for 3 hours. One representative experiment of n=3 independent experiments.

Finally, we wanted to identify which ubiquitin residues on STING are involved in its degradation. We predicted that ubiquitination at multiple lysine residues would drive STING degradation, since ESCRT sorting of cargo requires assembly of a heterotetrameric HGS/STAM complex containing up to 10 ubiquitin binding sites^50^. When we examined lysine conservation in human STING, we identified K20, K150, K236, K289, K338 and K370 as highly conserved, among 9 lysine residues (Fig. S9d). We excluded K289 from our experiments, since STING K289R is unstable and degraded at steady state (Fig. S9e-f), as previously reported^42^. We then generated 293Ts stably expressing a STING 3KR (K20R/K150R/K236R) and a STING 5KR (K20R/K150R/K236R/K338R/K370R) mutant fused to HA. When stimulated with cGAMP, STING 3KR showed a modest decrease in degradation, while STING 5KR almost completely abrogated it (Fig. 7d-e). STING 5KR was not sensitive to MLN7243 (Fig. S9g). We also tested whether STING 5KR could still exit the ER since, for example, K224 is critical for this process^42^. When stimulated with cGAMP, STING 5KR cells showed STING co-localization with GM130, suggesting that the defect in degradation was not due to defects in STING 5KR ER exit (Fig. S9h). A STING 2KR mutant (STING K338R/K370R) marginally reduced STING degradation (Fig. S9i-k), suggesting that K338 and K370 are not the only lysines driving STING degradation. To test if STING 5KR showed defects in association with ESCRT, we expressed STING 5KR-TurboID in 293T cells and performed pull-downs. STING 5KR failed to label HGS and VPS37A (Fig. 7f), suggesting that the defect in degradation is driven by defects in association with ESCRT.

Overall, these results suggest that STING ubiquitination is driven by STING oligomerization. Ubiquitination is triggered on multiple lysine residues driving STING association with VPS37A.

### A patient mutation in the ESCRT-I subunit UBAP1 induces a constitutive STING-dependent type I IFN response

We next investigated whether proteins in our dataset have established roles in human disease, potentially suggesting altered STING signaling as part of the underlying pathophysiology. Mutations in the ESCRT-I subunits VPS37A and UBAP1 have been shown to induce the neurodegenerative disease Hereditary Spastic Paraplegia (HSP)^51–54^. Interestingly, a subset of patients with mutations in the Aicardi-Goutiére Syndrome genes ADAR1, IFIH1 and RNASEH2B, that present with constitutive induction of type I IFN, have been shown to develop HSP^55^. Since UBAP1 is an endosome specific ESCRT subunit that does not play a role in cytokinetic abscission^25, 56, 57^, we asked if disease causing mutations in this gene could lead to dysfunctional STING degradation and exacerbate STING dependent signaling. Consistent with our mass spectrometry (Fig 1e, 3a), we found that STING-TurboID labeled UBAP1 (Fig. 8a). Pathogenic UBAP1 mutations in patients derive from frameshifts leading to truncated UBAP1 mutants containing only the N-terminal UBAP1-MVB12-associated (UMA) domain, responsible for interaction with ESCRT, and lacking the Solenoid of Overlapping Ubiquitin-Associated Domains (SOUBA), responsible for association with ubiquitinated cargo. These truncated mutants are dominant negatives for ESCRT function^56^. To mimic a pathogenic variant found in patients, we introduced a stop codon in place of residue G98 in UBAP1^51^ (UBAP1DN) (Fig. 8b) and fused it to mScarlet. When expressed in the reporter STING-HA cell line, UBAP1DN blocked STING degradation (Fig. 8c-e) consistent with our findings that a specific VPS37A/UBAP1 ESCRT complex regulates STING trafficking and degradation.

**Figure 8.**
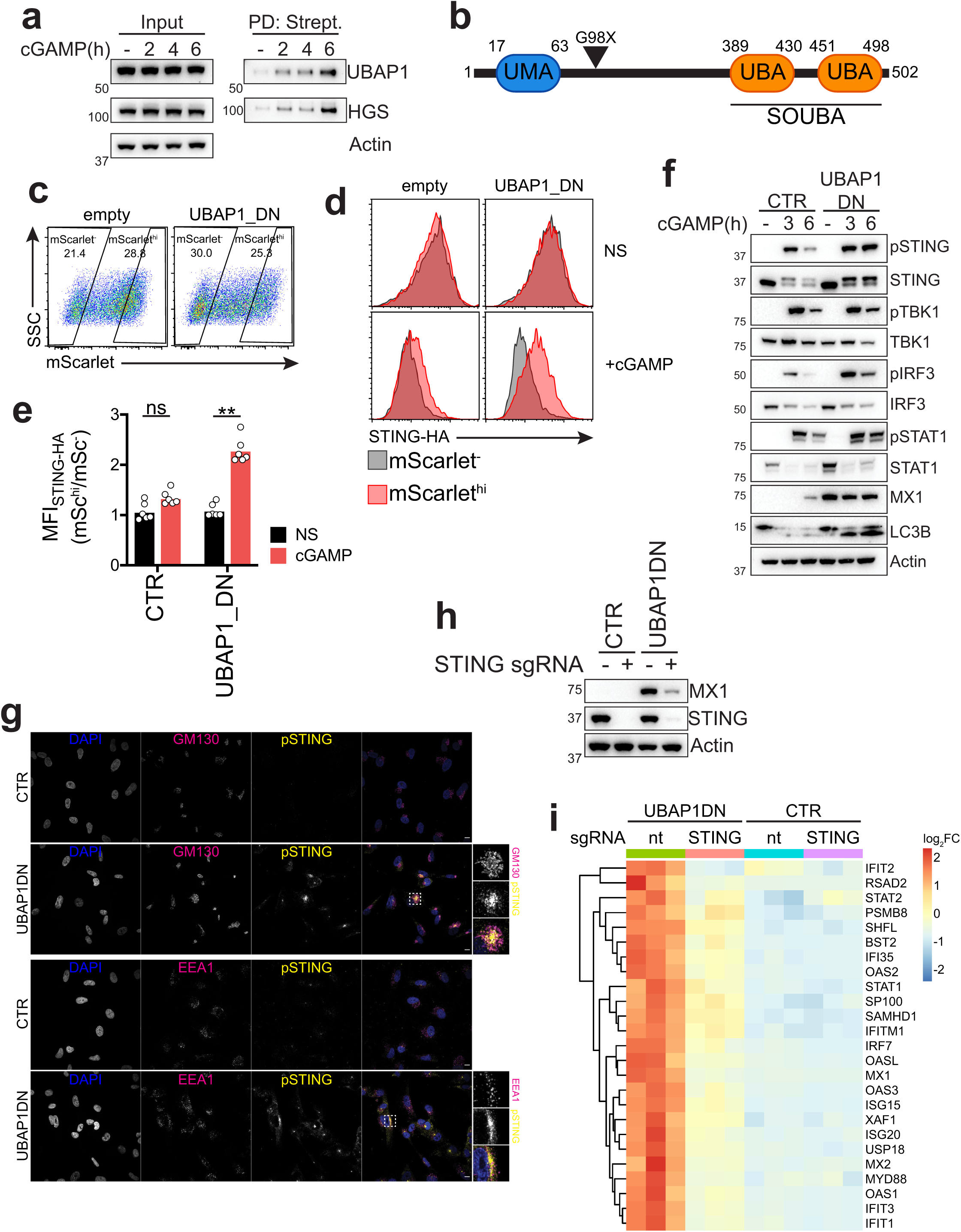
A patient mutation in the ESCRT-I subunit UBAP1 induces a constitutive STING-dependent type I IFN response. **a)** Immunoblot of the indicated proteins in the input or after streptavidin pulldown in 293T STING-TurboID stimulated with cGAMP for the indicated times. One representative blot of n=2 independent experiments. **b)** Scheme of UBAP1 domains and of the mutation introduced (stop codon at position G98) to obtain UBAP1DN. UMA: UBAP1-MVB12-Associated domain; SOUBA: Solenoid of ubiquitin associated domains (UBA). **c)** Expression of mScarlet-UBAP1DN after transfection in 293T stably expressing STING-HA and gating strategy for mScarlet^-^ and mScarlet^hi^ populations. **d)** STING-HA levels in cells as in c) in mScarlet^-^ (dark grey) and mScarlet^hi^ (red) that were either non-stimulated (NS) or treated with cGAMP for 6 hours. One representative plot of n=3 independent experiments with n=2 technical replicates per experiment. **e)** Median Fluorescence Intensity (MFI) of STING-HA signals shown as a ratio of MFI of the mScarlet^hi^ population over the MFI of the mScarlet^-^ population. n=3 independent experiments with n=2 technical replicates per experiment. Each dot represents an individual replicate. One-way ANOVA with Dunnet’s multiple comparisons. **p<0.01, ns: not-significant **f)** Immunoblot of the indicated proteins in BJ1 expressing mScarlet (CTR) or mScarlet-UBAP1DN stimulated with 2’3’-cGAMP for the indicated times. **g)** Immunofluorescence of DAPI (blue), GM130 or EEA1 (magenta) and phospho-STING (yellow) in BJ1 fibroblasts expressing either a control vector or UBAP1DN as in a). One representative field of n=5 field per condition for n=2 independent experiments. Scale bar is 10µm. **c)** Immunoblot of the indicated proteins. **h)** Immunoblot of the indicated proteins in BJ1 expressing mScarlet (CTR) or mScarlet-UBAP1DN transduced with spCas9 and a control sgRNA or a STING targeting sgRNA. One experiment representative of n=3 independent experiments. **i)** RNAseq derived heat-map for expression of selected ISGs in the indicated samples. Each column represents a technical replicate.

We then generated stable UBAP1DN expressing fibroblasts and stimulated them with cGAMP. UBAP1DN expressing fibroblasts showed a marked increase in STING signaling, reduction in STING degradation and accumulation of lipidated LC3B (Fig. 8f, S10a). Expression of the UBAP1DN mutant in primary human Monocyte Derived Dendritic Cells (MDDCs) also led to increased STING signaling (Fig. S10b). Interestingly, we noticed that UBAP1DN expressing fibroblasts induced the interferon-stimulated gene MX1 at steady state, in absence of cGAMP stimulation (Fig. 8f). The MX1 upregulation at steady state was also present in UBAP1DN expressing MDDCs, which are terminally differentiated cells and do not cycle, in addition to spontaneous DC maturation, shown by CD86 upregulation (Fig. S10b-d). This suggested that perturbation of ESCRT function by the expression of the HSP UBAP1 mutant, which blocks STING degradation, leads to a constitutive type I IFN response. Consistent with this hypothesis, KO of HGS and VPS37A in fibroblasts also led to MX1 upregulation at steady state (Fig. S10e).

Mutations in genes regulating STING trafficking, such as COPA, have been shown to induce STING dependent spontaneous type I IFN production due to accumulation of STING at the Golgi^22, 58–61^. Similar to mutations in COPA, we reasoned that activation of constitutive type I IFN in UBAP1DN expressing cells could be the result of impaired homeostatic STING trafficking, specifically post-Golgi, leading to intracellular accumulation of activated STING. Consistent with this hypothesis, UBAP1DN expressing fibroblast showed accumulation of phospho-STING partially co-localizing with the Golgi marker GM130 and the early endosome marker EEA1 (Fig. 8g). To test if accumulation of STING in the endolysosomal compartment is sufficient to spontaneously activate the sensor, we substituted the four transmembrane domains of STING with the four transmembrane domains of TMEM192 by fusing the STING C-terminal (aa139-379) to TMEM192 (TMEM192-STING). TMEM192 has been characterized as a membrane-resident protein of the late endosomal and lysosomal compartment^62^. Therefore, we hypothesized that the TMEM192-STING construct would be constantly trafficked to and accumulated post-Golgi, mimicking the UBAP1 mutant dependent block of STING trafficking. Expression of the TMEM192-STING construct led to dispersion of intracellular STING staining that partially co-localized with CD63 (Fig S10f). Compared to WT full length STING, TMEM192-STING led to spontaneous signaling activation, as shown by phospho-STING, even in absence of ligand (Fig. S10g).

To test if induction of ISGs in UBAP1DN expressing fibroblasts was due to accumulation of activated STING, we used CRISPR KO. When STING was deleted, UBAP1DN expressing fibroblasts showed a drastic reduction of MX1 induction (Fig. 8h, S10h). To confirm these results, we performed RNAseq. UBAP1DN expressing fibroblasts transduced with a non-targeting sgRNA showed an increase in ISG expression compared to control fibroblasts (Fig. 8i, S10i). GO analysis of differentially expressed genes identified genes involved in the “response to virus” or “type I IFN signaling” among the most upregulated in UBAP1DN expressing cells (Fig. S10i). STING KO in UBAP1DN expressing cells led to a specific reduction in the expression of ISGs compared to UBAP1DN expressing fibroblasts transduced with a non-targeting sgRNA, including genes involved in the type I IFN pathway as the most downregulated (Fig. 8i, S10i-j). Overall, these data suggest that STING is subject to a homeostatic degradative flux and perturbation of ESCRT leads to post-Golgi accumulation of STING which is sufficient for activation of the sensor.

cGAS has been shown to induce tonic ISG transcription at steady-state *in vitro* and *in vivo*, regardless of presence of exogenous DNA, suggesting that cGAS produces low levels of cGAMP in cells to prime this response^63, 64^. To test if cGAS was required to prime STING trafficking at steady state and contributed to the phenotype shown by pathogenic UBAP1 expressing cells, we knocked out cGAS in UBAP1DN expressing fibroblasts. cGAS KO reduced induction of the ISG MX1, suggesting that cGAMP production at steady state is responsible for priming the constitutive flux of STING trafficking (Fig. S10k). This is consistent with the phenotype shown in cells with COPA deficiency that activate a spontaneous type I IFN response that is both cGAS and STING dependent^58^.

Taken together, these data suggest that cells expressing a pathogenic mutant of UBAP1 have a heightened response to cGAMP through impaired ESCRT mediated STING degradation. Mutant UBAP1 leads to intracellular accumulation of post-Golgi activated STING with consequent constitutive type I IFN activation. Therefore, we propose that cGAS primes constant STING trafficking between the ER and the endosome for degradation at steady state, and functionality of an endosome specific VPS37A/UBAP1 containing ESCRT complex prevents STING accumulation preventing downstream signaling. Mutations in genes regulating STING trafficking could represent a general disease sensitizing mechanism leading to either a lowered threshold for STING activation or directly inducing STING dependent responses via disruption of the homeostatic STING degradative flux.

## Discussion

While some of the factors regulating STING exit from the ER have been characterized, the signals and genes regulating STING post-Golgi trafficking remained to be identified. To address this gap and unbiasedly identify genes involved in STING trafficking, we generated a time-resolved map of STING neighboring proteins at different intracellular compartments post cGAMP activation (Fig. 1) and carried out CRISPR screens for regulators of STING degradation (Fig. 2, 6), providing a basis for our studies and a resource for the field. By focusing on mechanisms of STING post-Golgi trafficking, we identified ubiquitin as the post-Golgi signal regulating STING degradation (Fig. 5). By performing targeted CRISPR screens, we identified UBE2N to drive STING polyubiquitination (Fig. 6), a process triggered by STING oligomerization (Fig. 7c). Similar to the signaling platform created at the TGN by phosphorylated STING for activation of IRF3 and induction of type I IFN, ubiquitinated STING in the endosomal compartment creates a platform for the recruitment of an endosome-specific VPS37A/UBAP1-containing ESCRT complex. Association with this complex drives degradation of the sensor via fusion of vesicles coated with oligomeric STING with the endolysosome, leading to reduction of lipidated LC3B. We therefore show that endosomal trafficking and autophagy resolution are both part of the same ESCRT coordinated signaling shutdown mechanism downstream of STING.

Based on our time-resolved map, we also show that expression of a UBAP1 mutant found in patients with hereditary spastic paraplegia leads to post-Golgi accumulation of activated STING with consequent constitutive induction of type I IFN (Fig. 8). Based on this evidence, we therefore propose an updated model of STING trafficking in which tonic cGAMP production primes a basal flux of STING trafficking from the ER to the lysosome with consequent ESCRT dependent constant degradation. Inactivating mutations in genes controlling STING trafficking represent a generalized mechanism for inducing constitutive STING-dependent responses that could lead to or exacerbate disease. A similar mechanism blocking steady-state TLR7 activation has been characterized, by which UNC93B1-Syntenin1 mediated trafficking of the receptor to MVBs is required for reduction of TLR7 intracellular levels and blunting of signaling at steady state^65^. Similarly, mutations in C9orf72 or NPC1, which block STING degradation at steady state, lead to spontaneous activation of the sensor^66, 67^. STING is highly conserved among organisms and we showed that accumulation of STING in the endosomal compartment is sufficient to induce STING responses (Fig. S10f-g). It is tempting to speculate that an ancestral function of STING in anti-pathogen defense could be sensing perturbation of intracellular trafficking pathways due to pathogen invasion. Interestingly, STING has been shown to be activated by HSV, influenza and HIV-1 viral entry independently of nucleic acids^68–70^. Further work will be necessary to link potential endosomal disruption to spontaneous activation of the sensor.

What is the trigger for STING dependent LC3B lipidation of membranes, what role the autophagy machinery plays in STING degradation and how it interacts with ESCRT remain open questions. LC3B lipidation upon STING activation has been shown to be driven by ATG16L1 dependent recruitment of ATPV1 subunits on single membranes^16, 71^. While we show that ESCRT regulates STING degradation, KO of ATG16L1 and ATG5 also reduces this process (Fig. S8b-f), suggesting a cross-talk between the two pathways. We propose that, while low tonic levels of cGAMP prime STING trafficking that can be removed by ESCRT at the endolysosome, presence of a high level of cGAMP in cells leads to marked STING oligomerization which in turn drives its UBE2N dependent ubiquitination and removal in a manner dependent both on ESCRT and the autophagy machinery (Fig. S11). LC3B presence on membranes has been shown to drive interactions with membrane fusion machineries such as the HOPS complex^72^, therefore ATG16L1 dependent lipidation of oligomerized STING containing vesicles would enhance degradation of STING oligomers. Identifying how LC3B lipidation of STING vesicles aids the degradative process and what membrane topology is resolved by ESCRT will help elucidate how the two processes interact.

Perturbation of STING trafficking is a potent tool to manipulate the pathway and regulate STING responses. While blocking STING degradation post-Golgi could be detrimental due to spontaneous activation of the sensor and lead to autoinflammation^58, 66, 67, 73^, precise and time-limited intervention could be desirable. Blocking lysosomal acidification with Bafilomycin A has been shown to improve STING dependent anti-tumor responses^17^, while blocking STING exit from the ER shuts down autoinflammation^74^. Our time-resolved map of STING trafficking provides the rationale to identify targets that could be perturbed in order to shut-down or increase the magnitude of STING responses.

## Authors contribution

M.G. and N.H. designed the study and wrote the manuscript. M.G. performed most of the experiments. B.L. helped with conceptualization and performed some experiments. M.P. and D.D. performed mass spectrometry and analysis. A.A. performed RNAseq of UBAP1DN expressing cells. M.A.S. performed sequencing and analysis of the ubiquitin screen and analyzed the RNAseq data. R.J.C. performed image quantification. J.G.D. and A.B. helped with the design of the CRISPR screens and analysis. S.C. helped with design of the proteomics experiments and editing of the manuscript. K.K. helped with statistical and computational analysis of the proteomics data.

## Acknowledgements

We thank the Flow Cytometry core at the Broad Institute for help with flow cytometry. We thank Mikolaj Slabicki, Matthew Bakalar and Lawrence D Schweitzer for discussions and Jon Kagan for feedback on the manuscript. The project was supported by NIH/NIAID U19 AI133524 (N.H.) and NIH/NHGRI P50HG006193 (N.H.). M.G. was supported by an EMBO Long-Term Fellowship (ALTF 486-2018) and is a Cancer Research Institute/Bristol Myers Squibb Fellow (CRI 2993). M.A.S. was supported by the National Cancer Institute of the National Institutes of Health under Award Number T32CA136432. R.J.C. is supported by a Fannie and John Hertz Foundation Fellowship and an NSF Graduate Research Fellowship. K.K. was supported by the National Cancer Institute (NCI) Clinical Proteomic Tumor Analysis Consortium through grant U24CA210979. The content is solely the responsibility of the authors and does not necessarily represent the official views of the National Institutes of Health.

## Methods

### Human Cell lines

293T (CRL-3216), hTert-BJ1 (BJ-5ta - CRL-4001) and U-937 (CRL-1593.2) were from ATCC. 293T were cultured in DMEM (Corning) supplemented with 10% FBS (VWR), 1X GlutaMax (Thermo Fisher) and 1X Penicillin/Streptomycin (Corning). hTert-BJ1 were cultured in a 4:1 mixture of DMEM (Corning):Medium 199 (Lonza) supplemented with 10% FBS (VWR), 1X GlutaMax (Thermo Fisher) and 1X Penicillin/Streptomycin (Corning). U-937 and THP-1 were cultured in RPMI (Thermo Fisher) supplemented with 10% FBS (VWR), 1X GlutaMax (Thermo Fisher) and 1X Penicillin/Streptomycin (Corning).

### Primary Human Cells

CD14^+^ monocytes were isolated from peripheral adult human blood as previously described^1^. CD14^+^ monocytes were cultured in RPMI (Gibco) supplemented with 10% FBS (VWR), 1X GlutaMax (Thermo Fisher), 50µg/ml Gentamicin (Thermo Fisher) and 1X Penicillin/Streptomycin (Corning). Human Monocyte Derived Dendritic Cells were differentiated as described in Gentili et al 2019^2^. Medium was replaced 1 days after transfection and cells were selected with 2µg/ml Puromycin (Invivogen). Cells were replated at 0.15×10^6c/w in a 96 well plate 5 days post differentiation, rested for 1 hour in the incubator and then stimulated with direct addition of cGAMP in the medium or with pI:C complexed with Lipofectamine.

### Constructs

The plasmids psPAX2 (#12260) and pCMV-VSV-G (#8454) were from Addgene. pSIV3+ was previously described^2^. pTRIP-hPGK-Blast-2A was cloned from pTRIP-CMV-STING-GFP (kind gift of Nicolas Manel) by Gibson assembly of PCR amplified hPGK from pCW57-MCS1-2A-MCS2 (Addgene #71782) and a gBlock (IDT) for Blasticidin resistance. pTRIP-UbC-Blast-2A-STING-mNeonGreen was cloned from Gibson assembly of PCR amplified UbC promoter from FUGW, PCR amplified Blast, PCR amplified STING and a gBlock for humanized mNeonGreen described in Tanida-Miyake et al 2018^3^. pTRIP-hPGK-STING-TurboID was cloned by Gibson assembly of PCR amplified TurboID from V5-TurboID-NES_pCDNA3 (Addgene #107169) and STING from pTRIP-CMV-STING-GFP. pTRIP-hPGK-Blast-2A-STING-HA/STING V155M-HA/STING R284S-HA/ were cloned by Gibson assembly from PCR amplification or PCR mutagenesis. pXPR101-Hygro was cloned by Gibson assembly of a gBlock for Hygromycin resistance into pXPR101 (kind gift of Broad GPP). CROPseq-guide-Puro was a kind gift of Paul Blainey. pXPR_BRD023 (lentiCRISPR v2) was a kind gift of the Broad GPP platform. sgRNAs were cloned by gateway cloning into the respective vectors of annealed primers. pTRIP-SFFV-mNeonGreen and pTRIP-SFFV-Blast-2A-STING-mNeonGreen were obtained by Gibson assembly. pTRIP-SFFV-Hygro-2A-mScarlet, pTRIP-SFFV-Hygro-2A-mScarlet-HGS and pTRIP-SFFV-Hygro-2A-mScarlet-VPS37A were obtained by Gibson assembly of PCR amplified Hygro from pXPR101-Hygro, PCR amplified HGS or VPS37A from U937 cDNA and PCR amplified mScarlet from pmScarlet_C1 (Addgene #85042) into pTRIP-SFFV-EGFP-NLS (Addgene #86677). pTRIP-SFFV-Hygro-2A-mScarlet-VPS4A E228Q was obtained by Gibson assembly of PCR amplified VPS4A E228Q from pEGFP-VPS4-E228Q (Addgene #80351). pTRIP-SFFV-Hygro-2A-mScarlet-UBAP1DN and pTRIP-SFFV-Puro-2A-mScarlet-UBAP1DN (truncated mutant at residue 97 - mutation G98X) were obtained by Gibson assembly of a gBlock for UBAP1DN. pTRIP-hPGK-Hygro-2A-FLAG-Ubiquitin and pTRIP-SFFV-Blast-2A-HA-Ubiquitin were obtained by Gibson assembly of gBlocks for FLAG-Ubiquitin or HA-Ubiquitin. pTRIP-hPGK-Blast-2A-TMEM192-STING-HA was obtained by TMEM192 PCR amplification from 293T cDNA, PCR amplification of STING (aa139-379) and Gibson assembly. pTRIP-hPGK-Blast-2A-STING 3KR-HA, pTRIP-hPGK-Blast-2A-STING 5KR-HA and pTRIP-hPGK-Blast-2A-STING 2KR-HA were obtained by Gibson assembly of gBlocks in pTRIP-hPGK-Blast-2A.

### Production of lentivirus and lentiviral transductions

Lentiviruses were produced as described in Gentili et al 2019^2^. Briefly, 0.8 million/well 293T in a 6 well plate were transfected with 1µg psPAX, 0.4µg pCMV-VSV-G and 1.6µg of viral genomic DNA with TransIT-293 (Mirus) and left O/N. To generate SIV-VLPs, cells were transfected with 2.6µg pSIV3+ and 0.4µg pCMV-VSV-G. Medium was then changed to 3ml of fresh medium corresponding to the cell line to be transduced. Supernatants were harvested 30-34 hours after medium changed and filtered at 0.45µm. 0.5 million 293T, hTert-BJ1 or U937 were infected with 2 ml of fresh virus in presence of 8µg/ml Protamine (Millipore Sigma). To generate transduced MDDCs, 2×10^6 freshly isolated CD14+ monocytes were transduced in a 6 well plate with 1ml of lentivirus and 1ml of SIV-VLPs with 8µg/ml protamine.

### STING-TurboID

#### Generation of cells and stimulation

293T cells were transduced with pTRIP-hPGK-Blast-2A-STING-TurboID and selected with 15µg/ml Blasticidin (Invivogen) for one week. To test the construct via pull-down, 0.8 million cells were seeded in a 6 well plate. The following day, cells were permeabilized with 300µl/well of cGAMP permeabilization buffer [50mM HEPES (Corning), 100mM KCl (Thermo Fisher), 3mM MgCl_2_ (Thermo Fisher), 0.1mM DTT (Thermo fisher), 85mM Sucrose (Thermo Fisher), 0.5mM ATP (Cayman chemicals), 0.1mM GTP (Cayman Chemicals), 0.2% BSA (Seracare), 0.001% Digitonin (Promega)] containing 1µg/ml 2’3’-cGAMP (Invivogen) or water for 10 minutes at 37°C, washed with 3ml of warm medium and then medium was replaced. For mass-spec, 6 million cells were seeded in a 10cm dish for each condition. Cells were stimulated with 2.8ml of cGAMP permeabilization buffer containing 6µg total of cGAMP per dish. Cells were left stimulating for the desired times and 500µM biotin (Cayman chemicals) was added in each well 30 minutes prior to harvest. Cells were harvested by trypsinization, washed 3 times in cold PBS and pellets were frozen until processing. For experiments in 6 well plates, 3 wells per condition were harvested. For experiments in 10cm dishes, one dish per condition was harvested.

#### Pull-down

Cells were lysed on ice in 550µl (6 well plates) or 1ml (10cm dishes) of RIPA buffer (Boston Bioproducts) in presence of cOmplete, Mini, EDTA-free Protease Inhibitor Cocktail (Millipore Sigma) and PhosSTOP (Millipore Sigma) for 10 minutes. Lysates were cleared by centrifugation at 16000g for 10 minutes at 4°C. 10% of the lysed cells was set aside as input. set aside as input. Pull-down and washes were performed as in ^4^. Briefly, lysates were mixed with Pierce Streptavidin Magnetic Beads (Thermo Fisher) at a ratio of 100µl beads/4 million cells. Lysates were incubated with beads with constant rotation for one hour at room temperature and then overnight (O/N) at 4°C. Beads were then applied to a magnet and subjected to the following washes: 2 times with 1ml of RIPA, 1 time with 1M 1ml of KCl (Thermo Fisher), 1 time with 1ml of 0.1M Na_2_CO_3_ (VWR), 1 time with 1ml of freshly prepared 2M Urea (VWR) resuspended in 10mM Tris-HCl pH 8.0 (Thermo Fisher) and 2 times with RIPA. Proteins were eluted from beads by adding 150µl (6 well plates) or 500µl (10cm dish) of non-reducing Laemmli (Boston bioproducts) containing 20mM DTT (Thermo Fisher) and 2mM biotin (Cayman chemicals) and boiled for 20 minutes. Input was diluted with 2X sample buffer (Sigma). For mass spectrometry, beads were processed as follows.

#### Sample Processing for mass spectrometry

Co-IP was performed using 2.2mg of HEK293T cells expressing hPGK-Blasticidin-P2A-STING TurboID, 200 uL of Streptavidin beads, in duplicates, at 5 time points: not-stimulated, 30 minutes, 1, 2 and 6 hours.

Samples were received in duplicates, each in 1mL RIPA lysis buffer. Beads were washed with 50 mM Tris HCL (200 uL, pH 7.5, 2X) and transferred to fresh 1.5 mL eppendorf tubes. Beads were further washed with 2 M Urea/50 mM Tris HCL (200 uL, pH 7.5, 2X). Proteins were digested with trypsin (5 ug/mL, 80 uL) in 2 M urea/50 mM Tris HCL/1 mM dithiothreitol (DTT)) at 25 ℃ for 1 hr). Following a brief centrifugation step using a table-top centrifuge (5-10s), supernatants were transferred to clean 1.5 mL eppendorf tubes. Beads were washed once with 2 M urea/50 mM Tris HCL (60uL, pH 7.5, 2X) and supernatants were combined with respective supernatants from the first centrifugation step. Combined supernatants were centrifuged at 5000 g for 30s to pellet remaining beads and the supernatants were transferred to clean 1.5 mL eppendorf tubes.

Samples were reduced with DTT (4 mM) using a shaker (1000 rpm) for 30 min at 25°C) and alkylated with iodoacetamide (IAA, 10 mM) for 45 min at 25℃ in the dark. Proteins were digested overnight with trypsin (0.5 ug in trypsin buffer) at 25℃ using a shaker (700 rpm). Samples were acidified with formic acid (FA, 1%, 200 uL pH <3) and peptides were desalted using C18 stage tips (2 punches) following standard protocol ^5^. Briefly, stage tips were activated with 50% ACN, 0.1% FA (50uL, 1500 rcf) and conditioned with 0.1% FA (50 uL, 1500 rcf, 2X). Samples (350uL) were loaded on the tips and spun at 1500 rcf until all volume flowed through completely without drying the stage tips. Samples were washed with 0.1% FA (50uL, 2X, 1500 rcf), eluted with 50%ACN/0.1% FA (50 uL, 1500 rcf) and lyophilized. Peptides were subsequently reconstituted in fresh HEPES (50 mM, 95.3 uL, pH 7.5) for TMT labeling. Samples were labeled with TMT as follows: Non-Stimulated (126, 129N), 0.5 hr (127N, 129C), 1h (127C, 130N), 2h (128N, 130C), 6 hr (128C, 131). TMT labeling (240 ug per sample) occurred for 1 hr at room temperature with shaking (800 rpm), following standard protocol. Samples were quenched with 5% hydroxylamine (8 uL, 20℃, 700 rpm), all channels were combined in one vial and lyophilized. The combined samples were reconstituted in 0.5% acetic acid (100 uL) and fractionated following standard protocol ^6^. Briefly, 3 SCX discs (polytetrafluoroethylene (PTFE) material) were placed in 200 uL pipette tips followed by 2xC18 discs on top. Tips were conditioned with methanol (100 uL, 3500g, 1 min), followed by 0.5% acetic acid/80% ACN (100 uL, 3500 g, 1 min) and 0.5% acetic acid (100 uL, 3500 g, 1 min). Tips were equilibrated with 0.5% acetic acid (100 uL, 3500 g, 1 min), 500 mM NH_4_AcO/20% ACN (100 uL, 3500 g, 1 min) and 0.5% acetic acid (100 uL, 3500 g, 1 min) prior to sample loading (100 ul, 3500 g). The sample was washed twice with 0.5% acetic acid (100 uL, 3500 g, 1 min, 2X), followed by 0.5% acetic acid/80% ACN (100 uL, 3500g, 1 min). A stepwise elution occurred using 50 mM NH_4_AcO/20% ACN (pH 5.15, 50 ul, 3500 g, 1 min, fraction 1), 50 mM NH_4_HCO_3_/20% ACN (pH 8.25, 50 ul, 3500 g, 1 min, fraction 2) and 0.1% NH_4_OH/20% ACN (pH 10.3, 50ul, 3500g, 1 min, fraction 3). Acetic acid (0.5%, 200ul) was added to each eluate to reduce ACN concentration to <5%. Fractions were subsequently desalted using 2 punch C18 stage tips following the protocol described above and eluted with 80%ACN/0.5% acetic acid (60 uL, 1500 rcf). Samples were lyophilized and re-suspended in 3%ACN/0.1%FA (10 ul) for nanoLC-MS/MS analysis.

#### MS Analysis

Fractionated samples were analyzed on an Orbitrap Q-Exactive HF Plus MS (Thermo Fisher Scientific) equipped with a nanoflow ionization source and coupled to a nanoflow Proxeon EASY-nLC 1000 UHPLC system (Thermo Fisher Scientific). Acquisition occurred in positive ion mode. Samples were injected on an in-house packed column (22cm x 75um diameter C18 silica picofrit capillary column) heated at 50℃. The mobile phase flow rate was 250 nL/min of 3% ACN/1% FA (solvent A) and 90% ACN/ 0.1% FA (solvent B). Peptides were separated using the following LC gradient: 0-6% B in 1 min, 6-30% B in 85 min, 30-60% B in 9 min, 60-90% B in 1 min, stay at 90% B for 5 min, 90-50% B in 1 min, and stay at 50% B for 5 min. Data was acquired in centroid mode for both MS1 and MS2 scans. Samples were analyzed in data dependent analysis (DDA) mode using a Top-12 method. Ion source parameters were: spray voltage 2 kV, source temperature 250 °C. Full MS scans were acquired in the m/z range 200–2000, with an AGC target 3e6, maximum IT 10 ms and resolution 70,000 (at m/z 200). MS/MS parameters were as follows: AGC target 1e5, maximum IT 50 ms, loop count 10, isolation window 1.6 m/z, isolation offset 0.3 m/z, NCE 31, resolution 17,500 (at m/z 200) and fixed first mass 100 m/z; unassigned and singly charged ions were excluded from MS/MS.

#### Proteomic Data Analysis

Raw MS data were analyzed using Spectrum Mill Proteomics Workbench (prerelease version B.06.01.202, Agilent Technologies). A trypsin-specific enzyme search was performed against 2017 uniprot human fasta file (UniProt.human.20171228.RISnrNF.553smORFs.264contams) containing 65095 entries. Peptide and fragment tolerances were at 20 ppm, minimum matched peak intensity 40% and peptide false discovery rates (FDR) were calculated to be <1% using the target-decoy approach^7^. Fixed modifications were carbamidomethylation, TMT 10 (N-term, K) and variable modifications were Acetyl (ProN-term), Oxidized methionine (M), Pyroglutamic acid (N-termQ) and Deamidation (N). Spectra with a score <4 were filtered out. Peptides were validated using the following parameters: for charge states 2-4, a FDR of 1.2 was applied to each run and for charge state 5, a FDR of 0.6 was applied across all runs. Results were further validated at the protein level and proteins with a score of 20 or higher were accepted as valid. Reporter ion correction factors, specific to the TMT batch, were applied. A protein/peptide summary was generated using the median across all TMT channels as the denominator. Shared peptides were assigned to the protein with the highest score (SGT).

#### Bioinformatics Analysis

Calculated ratios at the protein level were imported into Protigy v0.8.X.X for normalization and features selection (https://github.com/broadinstitute/protigy). To account for variability between samples, log ratios were normalized by centering using the sample Median and scaled using the sample Median Absolute Deviation (Median MAD). A moderated F-test^8^ was performed to identify proteins whose expression changed significantly across time points. Temporal profiles of significant proteins (FDR p-value <0.05) were z-scored and further subjected to fuzzy c-means clustering implemented in the e1071 R package. The number of clusters was set to three upon visual inspection of temporal profiles. The optimal fuzzification parameter m was determined as described in^9^. Gene Ontology (GO) overrepresentation analysis of proteins in the resulting clusters was performed with the gProfiler R-package^10^.

#### Network representation and analysis

The filtered dataset on adj p.value <0.07 was uploaded on STRING to generate a network. The network was then imported in Cytoscape (v3.8.1) and analyzed with stringApp (v1.6.0). Subcellular compartments were assigned by filtration in stringApp. Maps representing enrichments at different time-points were plotted by using the Hierarchical Clustering function in clusterMaker.

#### GO and Reactome enrichments

GO and Reactome enrichments present in Fig. S3 were calculated using the Functional Enrichment Analysis with values/ranks in STRING^11^.

### Genome-wide CRISPR screen

#### Generation of cells and screen

293T were transduced with pTRIP-UbC-Blast-2A-STING-mNeonGreen(mNG) and selected with 15µg/ml of Blasticidin (Invivogen) for one week. mNG^hi^ cells were sorted on a Sony SH-800. Cells were then transduced with pXPR101-Hygro to introduce spCas9, and kept under selection with 320µg/ml Hygromicin for the time of culturing. Cells were then transduced with the human targeting genome-wide sgRNA library Brunello^12^ at MOI 0.3 at a 1000X coverage (80 million cells). The library in lentiviral form was obtained from the Broad GPP. Cells were then selected in 2µg/ml Puromycin and passaged maintaining 1000X representation of the library for one week. The day prior to the stimulation, cells were plated at 20 million/T225. The cells were then stimulated by adding 40 ml of fresh medium containing 1µg/ml of 2’3’-cGAM(PS)2 (Invivogen) for 24 hours. Cells were then lifted, resuspended in MACS buffer (0.5% BSA, 2mM EDTA in PBS) and sorted on two Sony SH-800 at a 4000X coverage (320 million total cells sorted). Cells were then pelleted, washed with PBS, and pellets were frozen until DNA extraction. DNA was extracted with DNeasy Blood & Tissue Kit (Qiagen) following manufacturer’s recommendations.

#### Sequencing and screen analysis

Extracted DNA was submitted to the Broad Genetic Perturbation Platform (GPP) for Next Generation Sequencing. After deconvolution, reads per barcode were analyzed with the GPP Pooled Screen Analysis Tool using the Hypergeometric method (https://portals.broadinstitute.org/gpp/public/analysis-tools/crispr-gene-scoring).

#### Screen validation

293T STING-mNG spCas9 were transduced with SEC24C or ATP6V1G1 sgRNAs cloned in CROPseq-Guide-Puro and selected with 2µg/ml Puromycin (Invivogen) for one week. 0.016 million cells/well were plated in a 96 well plate the day prior to the stimulation, and then stimulated with 100µl of fresh medium containing 2’3’-cGAM(PS)2 (Invivogen) for 24 hours. For HGS and VPS37A, 293T STING-mNG cells were transduced with pXPR023 (lentiCRISPR v2) expressing sgRNAs for each of the genes and selected on Puromycin for one week. 0.125 million cells/well were plated in a 24 well plate the day prior to stimulation, and then stimulated with 500µl of fresh medium containing 2’3’-cGAM(PS)2 (Invivogen) for 24 hours.

### Ubiquitin targeted CRISPR screen

The library contained guides targeting 669 E3 and adaptors (compiled from Medvar et al., 2016^13^ and Li et al., 2008^14^), 40 E2 from Interpro, 7 E1, 28 Autophagy core proteins and 10 positive controls from the genome-wide CRISPR screen and was synthetized and cloned by the Broad GPP. Cells were generated as for the genome wide CRISPR screen. STING-mNG cells were sorted without fixation while STING-HA cells were sorted after fixation and staining as described in the flow cytometry paragraph. Both cell lines were sorted in 4 bins (top 5%, second top 5%, bottom 5%, second bottom 5%) at 4000X coverage. For STING-mNG DNA was extracted as for the genome-wide CRISPR screen. For STING-HA DNA was extracted with Quick-DNA FFPE kit (Zymo). Sequencing was performed as described in Fulco et al 2016^15^. Analysis was performed as for the genome-wide CRISPR screen.

### Co-Immunoprecipitation in 293Ts

293T were transduced with either pTRIP-hPGK-Blast-2A, pTRIP-hPGK-Blast-2A-STING-HA, pTRIP-hPGK-Blast-2A-STING V155M-HA or pTRIP-hPGK-Blast-2A-STING R284S-HA and selected with 15µg/ml Blasticidin for one week. Cells were then plated at 0.8 million cells/well in a 6 well plate and transfected with either pTRIP-SFFV-Hygro-2A-mScarlet-HGS or pTRIP-SFFV-Hygro-2A-mScarlet-VPS37A with TransIT-293 (Mirus) (3µg DNA/well). 24 hours post-transfection, 3 wells per condition were harvested via trypsinization. Cells were washed with PBS and lysed 550µl of Co-IP buffer (20 mM Tris-HCl pH 7.5, 150 mM NaCl, 0.5% NP-40 on ice for 30 minutes and cleared by centrifugation at 16000g for 20 minutes). 10% of the lysate was saved as input. The lysates were then incubated with Pierce Anti-HA Magnetic Beads (Thermo Fisher) at a concentration of 100µl beads/4 million cells O/N at 4°C. Beads were washed 5 times with Co-IP buffer and proteins were eluted by adding 150µl of non-reducing Laemmli (Boston bioproducts) containing 20mM DTT (Thermo Fisher) and boiled for 20 minutes. Input was diluted with 2X sample buffer (Sigma).

### Flag-Ubiquitin immunoprecipitation

293T cells were transduced with pTRIP-hPGK-Blast-2A-STING-HA and pTRIP-hPGK-Hygro-2A-FLAG-Ubiquitin and selected with 15µl Blasticidin (Invivogen) and 320µg/ml Hygromycin (Invivogen) for one week. 1 million cells/well were plated in a 6 well plate the day prior to the stimulation. Cells were then stimulated with 300µl cGAMP permeabilization buffer containing 1µg/ml cGAMP for 10 minutes, washed with 3ml of warm medium, and the medium was replaced. 3 wells per condition were harvested 2 hours post stimulation, washed with PBS and lysed with 550µl of RIPA buffer for 10 minutes on ice. Lysates were cleared at 16000g for 10 minutes at 4°C. 10% of the lysate was saved as input. The remaining lysates were incubated with 150µl

Pierce Anti-DYKDDDDK Magnetic beads (Thermo Fisher) O/N at 4°C with constant rotation. Beads were washed 3 times with a buffer containing 10mM Tris-Hcl pH7.5 (Thermo Fisher), 2mM EDTA (Thermo Fisher), 1% Nonidet-P40 Substitute (Roche) and 50mM NaCl (Thermo Fisher), and 2 times with RIPA buffer. Proteins were eluted by adding 150µl of non-reducing Laemmli (Boston bioproducts) containing 20mM DTT (Thermo Fisher) and boiled for 20 minutes. Input was diluted with 2X sample buffer (Sigma).

### HA-Ubiquitin immunoprecipitation in THP-1

5 million THP-1 expressing HA-Ubiquin (from pTRIP-SFFV-Blast-2A-HA-Ubiquitin) were plated in 5ml of fresh medium containing 5µg/ml digitonin (Promega) per well in a 6 well plate the day of the stimulation. Cells were pre-treated with 1µM H-151 for 30 minutes and then cGAMP was added to the well at a final concentration of 10µg/ml. Cells were stimulated for 4 hours, pelleted, washed with PBS and pellets were frozen at -80°C until immunoprecipitation. Pellets were lysed on ice for 15 minutes in 770µl Ubiquitin IP buffer [50mM Tris HCl pH7.5 (Corning), 150mM NaCl, 1mM EDTA (Gibco), 0.2% NP-40 (Millipore Sigma)] supplemented with cOmplete Mini EDTA-free Protease Inhibitor Cocktail (Millipore Sigma), PhosSTOP (Millipore Sigma) and 50µg/ml PR-619 (Lifesensors). Lysates were cleared by centrifugation at 16000g for 10 minutes at 4°C and 70µl were recovered directly in Sample Buffer 2X (Millipore Sigma) as input. The remaining 700µl were loaded on 75µl Pierce Anti-HA Magnetic Beads (Thermo Fisher) and pulled down at 4°C for 3 hours with constant rotation. Beads were then washed 5 times with Ubiquitin IP buffer and proteins were eluted in Laemmli SDS Sample Buffer 1X with beta-mercaptoethanol (Boston Bioproducts) by boiling at 95°C for 10 minutes. Magnetic beads were discarded and samples were subjected to western blot analysis.

### U937 and hTert-BJ1 stimulation for Western blotting

To obtain KO U937 and hTert-BJ1 for HGS and VPS37A, cells were transduced with pXPR023 expressing the corresponding guides and selected with 2µg/ml Puromycin for one week. In regard to HGS, U937 were transduced with HGS_g01 and HGS_g02, while hTert-BJ1 were transduced with HGS_g01 and HGS_g04. To stimulate U937, 0.2 million cells/well were seeded in a 96 well plate U bottom in 100µl and stimulated by adding 100µl of fresh medium containing cGAMP to a final concentration of 20µg/ml for 6 hours. Two wells per condition were harvested 6 hours post-stimulation, washed with PBS and pellets were frozen. For hTert-BJ1, 0.25 million cells/well were seeded in a 6 well plate the day before stimulation. Cells were then stimulated with 300µl of cGAMP permeabilization buffer containing cGAMP at 0.5µg/ml for 10 minutes, washed with 3ml of warm medium, and then medium was replaced. Cells were harvested at the indicated time-points post stimulation, washed with PBS, and pellets were frozen.

### U937 stimulation for cell death

0.2×10^6 cells per well were plated in a 96w plate U bottom and stimulated with direct addition of 20µg/ml cGAMP in the medium. Cells were stimulated for 24h. Half the cells were then recovered and used for Cell Titer Glo assay (Promega). Half the cells were stained with Annexin V Apoptosis Detection Kit (Biolegend) following manufacturer’s instructions.

### Dominant negative transfections for STING degradation

293T cells were transduced with pTRIP-hPGK-Blast-2A-STING-HA and selected with 15µg/ml Blasticidin (Invivogen) for one week. 0.08 million cells/well were seeded in a 24 well plate and transfected with TransIT-293 (Mirus) with 0.5µg/well of either pTRIP-SFFV-Hygro-2A-mScarlet, pTRIP-SFFV-Hygro-2A-mScarlet-VPS4A E228Q or pTRIP-SFFV-Hygro-2A-mScarlet-UBAP1DN and medium was replaced after O/N incubation. Cells were stimulated 40 hours post-transfection with 200µl/well of cGAMP permeabilization buffer containing 2µg/ml 2’3’-cGAMP for 10 minutes. Cells were then washed with 2ml of medium and medium was replaced. Cells were lifted and stained as indicated in the Flow Cytometry paragraph.

### Treatments with drugs

MLN7243 (Selleckchem) was used at 0.5µM in all experiments. All cell lines were pre-treated for one hour before cGAMP stimulation. 293T STING-mNeonGreen were plated the day before stimulation in a 24w plate at 0.2 million cells/well and were stimulated by adding 4µg/ml 2’3’-cGAM(PS) in the medium for 6 hours. hTert-BJ1 were seeded the day before stimulation in a 6 well plate at 0.25 million cells/wells. Cells were stimulated with 300µl cGAMP permeabilization buffer containing 0.5µg/ml 2’3’-cGAMP for 10 minutes, washed with 3ml of medium, and medium was then replaced. MLN7243 was added again after medium replacement. Cells were stimulated for the indicated times. Bortezomib final concentration was 1µM, MG-132 2µM, BafA1 100nM, H-151 0.5µM. For stimulation of CD14+ monocytes, 0.2 million cells per well were plated in a 96 well plate U bottom in 200µl. Cells were pre-stimulated with 0.5µM MLN7243 and then cGAMP was added directly to the well at a final concentration of 5µg/ml. Cells were stimulated for 8 hours.

### Flow cytometry

For flow cytometry analysis of 293T STING-mNG, cells were lifted with TrypLE (Thermo Fisher), washed in medium and resuspended in FACS buffer (1% BSA, 1mM EDTA, 0.01% NaN_3_ in PBS). For experiments involving intracellular staining of HA, 293T expressing HA-tagged WT or STING mutants, cells were lifted with TrypLE (Thermo Fisher), washed with PBS and stained using BD Cytofix/Cytoperm (BD Biosciences). Cells were fixed in Cytofix for 1 hours, washed twice with cytoperm, and stained with

Alexa Fluor 647 anti-HA.11 Epitope Tag Antibody (BioLegend) for one hour. Cells were then washed twice with Cytoperm and resuspended in FACS buffer for flow cytometry. For Propidium Iodide staining of CD14+ monocytes stimulated with cGAMP and MLN7243, cells were stained with Propidium Iodide Solution (Biolegened) following manufacturer’s recommendations. Acquisition was performed on a Cytoflex S or Cytoflex LX (Beckman Coulter). Data was analyzed with FlowJo (BD).

### Immunofluorescence

293T STING-TurboID were seeded directly on coverslips. 293T STING-HA were seeded on Fibronectin bovine plasma (Sigma - stock: 100X) coated coverslips. Cells were seeded the day before stimulation at 0.1 million cells/well density in 24 well plates. Cells were stimulated with cGAMP permeabilization buffer containing 1µg/ml cGAMP for 10 minutes, washed with warm medium, and incubated for the indicated times. hTert-BJ1 mScarlet or mScarlet-UBAP1DN were seeded Fibronectin bovine plasma coated coverslips at 0.05 million cells/well density in a 24 well plate and fixed 6 hours post seeding. Cells were then fixed with 2% Paraformaldehyde (Electron Microscopy Sciences) in PHEM buffer (Electron Microscopy Sciences) for 30 minutes at 37°C, washed three times with PBS and quenched with freshly prepared 0.1M Glycine for 10 minutes. Coverslips were then permeabilized and blocked with 10% goat serum (Thermo Fisher) in PBS, 0.5% BSA (Seracare), 0.05% Saponin from quillaja barka (Sigma) for 30 minutes. Coverslips were then stained with primary antibodies for 1-2 hours at room temperature in PBS, 0.5% BSA (Seracare), 0.05% Saponin from quillaja barka (Sigma) with 10% goat serum, washed 5 times, and then stained with secondary antibodies in PBS, 0.5% BSA (Seracare), 0.05% Saponin from quillaja barka (Sigma) for 1-2 hours. Coverslips were then washed 5 times, mounted with Fuoromont-G, with DAPI (Thermo Fisher) and dried at 37°C for one hour. Images were acquired on an Olympus IX83 using an Olympus PlanApo N 60X 1.42NA oil immersion objective controlled by Fluoview software. Images were analyzed in FiJi. TMEM192-STING images in Fig. S9 were acquired on a Ti-2 Eclipse inverted epifluorescence microscope (Nikon) using a 20X 0.75 NA CFI Plan Apo λ objective (Nikon).

### Image quantification

CellProfiler version 4.2.1^16^ was used to extract image-based features from 3-5 fields of view and two separate experiments per condition. Background subtraction was first performed on all channels and cross-channel colocalization values were measured using the MeasureColocalization module, including only pixels above 15% intensity for Overlap, Manders, and RWC measurements. Puncta were identified using the IdentifyPrimaryObjects module using global Minimum Cross-Entropy thresholding.

### Western blotting

Samples for pull-downs or immunoprecipitations were treated as described in the corresponding paragraphs. For experiments involving seeding of cells in 6 well plates or 24 well plates (293Ts and hTert-BJ1), one well per condition was harvested, lysed in RIPA buffer (Boston Bioproducts) containing cOmplete, Mini, EDTA-free Protease Inhibitor Cocktail (Millipore Sigma) and PhosSTOP (Millipore Sigma) for 10 minutes on ice. Lysates were cleared by centrifugation at 16000g for 10 minutes at 4°C and Laemmli 6X, Sample buffer, SDS, Reducing (Boston Bioproducts) was added prior to loading. Samples were run on NuPAGE 4 to 12%, Bis-Tris Gels (Thermo Fisher) and transferred on nitrocellulose membrane with an iBlot2 (Thermo Fisher). Membranes were blocked in 5% non-fat milk in TBS Tween. Antibodies against phospho proteins were incubated in 5% BSA TBS tween. ECL signal was recorded on a ChemiDoc Biorad Imager. Data was analyzed with ImageLab (Biorad).

### RT-qPCR

BJ1 fibroblasts were stimulated as described in the “U937 and hTert-BJ1 stimulation for Western blotting” section. Cells were harvested and pellet was frozen at -80°C until processing. RNA was extracted using the Quick-RNA Microprep kit (Zymo) following manufacturer’s instructions. 5µl of RNA (roughly corresponding to 1µg) was reverse transcribed to cDNA using the LunaScript RT SuperMix kit - dye based qPCR detection (NEB) following manufacturer’s instructions. 1µl of the reverse transcribed cDNA was used in each reaction for qPCR using the Luna Universal qPCR Master Mix (NEB) in a 10µl final reaction in a 384 well plate. 3 technical replicates per sample were measured on a CFX384 Touch Real-Time PCR Detection System (Biorad). Cycles were as follows: 1. Initial denaturation - 95°C, 60s; 2. Denaturation - 95°C, 15s; 3. Extension - 60°C, 30s; Melt Curve - 60-95°C. Steps 2-3 were repeated for 40 cycles. Primers were: IFNB1 FWD - CAGCATCTGCTGGTTGAAGA, RV - CATTACCTGAAGGCCAAGGA; IL6 FWD - CCCCTGACCCAACCACAAAT, RV - ATTTGCCGAAGAGCCCTCAG; GAPDH FWD - GTCTCCTCTGACTTCAACAGCG, RV - ACCACCCTGTTGCTGTAGCCAA. Data was analyzed using the 2^-ΔCt^ method over GAPDH.

### RNAseq

#### RNA Isolation

0.25 million cells/well in a 6 well plate were plated the day before harvesting in triplicates. Cells were left in culture for 24 hours before harvesting. RNA was isolated using the AllPrep DNA/RNA Mini Kit (Qiagen# 80204). Following total RNA extraction, mRNA was purified using Dynabeads^®^ Oligo(dT)25 (ThermoFisher# 61005).

#### Generation of RNA-seq libraries

Bulk RNA-seq libraries were assembled from purified mRNA using the Smart-Seq3 workflow^17^. The reactions were scaled 8-fold per sample, using 15ng input mRNA. First-strand synthesis and template switch reactions were performed by combining RT mix 1 and RT mix 2 and incubated in the thermocycler conditions noted in table First-strand synthesis. cDNA amplification was performed using 20µL of the first-strand synthesis reaction as input in the cDNA PCR mix and thermocycled as noted in table cDNA amplification. Amplified cDNA was cleaned up using 0.8x concentration of Ampure XP beads (Beckmen Coulter #A63881). Following clean up, cDNA samples underwent tagmentation and subsequent final library amplification. Briefly, 4ul of tagmentation mix was combined with 400pg of cDNA sample diluted in 4ul of H2O. The tagmentation reaction was incubated at 55 °C for 10min, with reaction being stopped with addition of 2µL 0.2%SDS. Tagmented samples were then added to the final library mix and amplified in the conditions noted in table Final library amplification. Final libraries were cleaned using 0.8x concentration of Ampure XP beads (Beckmen Coulter #A63881) and quantified using the Agilent Bioanalyzer High Sensitivity DNA (Agilent# 5067-4626) system.

#### RNA-Seq Analysis

Libraries were sequenced on an Illumina NextSeq 500 with paired-end reads at a depth of 10-15 million reads per sample. Reads were extracted and demultiplexed using bcl2fastq2 (v. 2.20.0) and sequencing quality was assessed with FastQC (v. 0.11.9), after which adapter and quality filtering was performed with Cutadapt (v. 3.1)^18^. Reads were mapped with Salmon (v. 1.4.0)^19^ using a whole genome decoy-aware transcriptome index built from GENCODE GRCh38 release 36, and mapping rates for all samples were between 90-93%. Differential expression analysis was performed in R with DESeq2 using a standard workflow. Our criterion for identifying DEGs was padj ≤ 0.01. Volcano plots were plotted with VolcaNoseR^20^. GO enrichment plots were calculated using the Functional Enrichment Analysis in STRING.

## Data analysis and statistical tests

Unless otherwise specified, data was analyzed and statistics were calculated using Graphpad PRISM (v9).

## Data availability

The datasets generated during and/or analyzed during the current study are available from the corresponding author on reasonable request. RNA-seq data will be deposited at NCBI GEO.

## List of sgRNAs

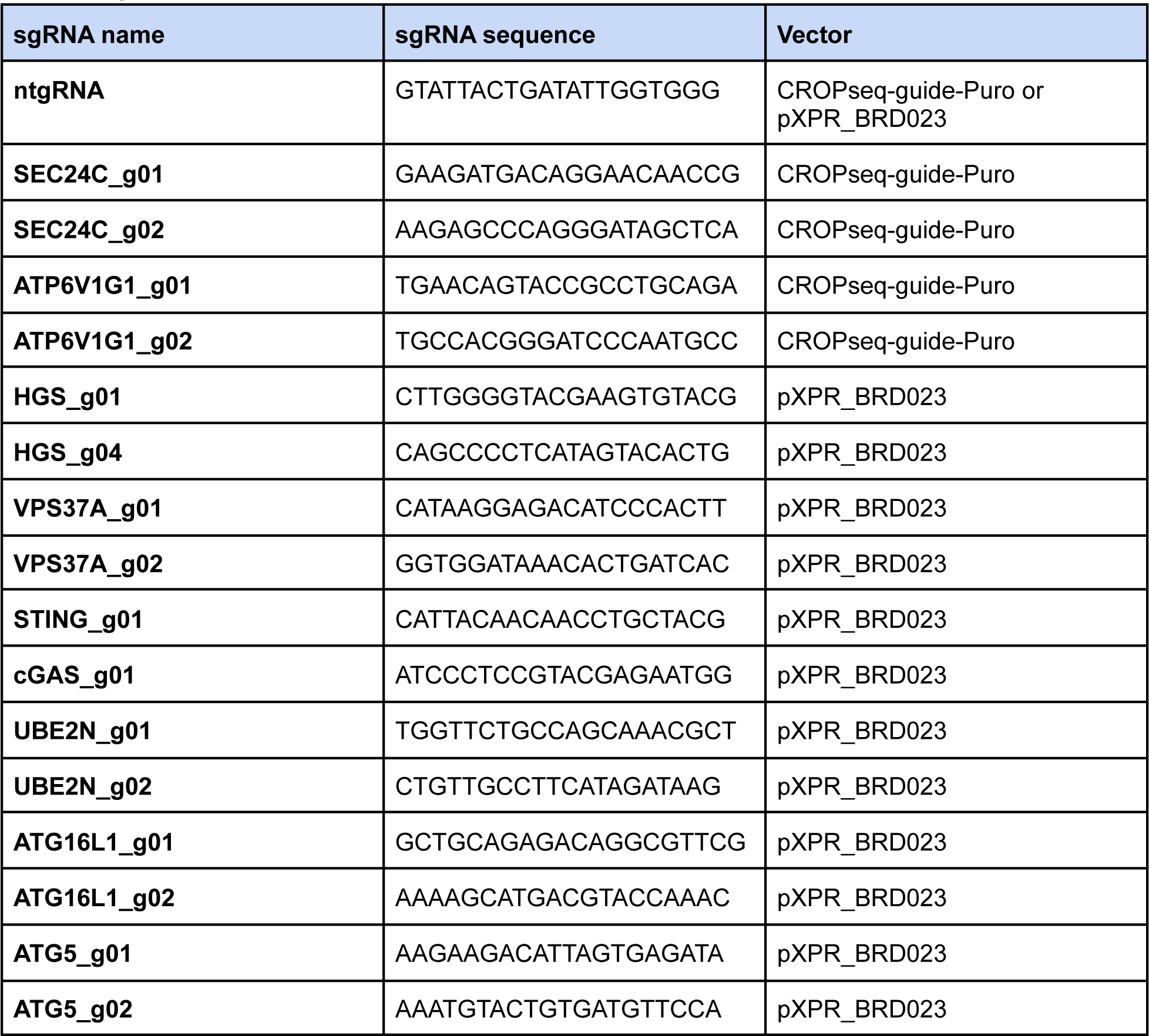

## List of antibodies

| Target | Type | Application | Source | Reference |
| --- | --- | --- | --- | --- |
| STING (D2P2F) | Rbt monoclonal | WB and IF | CST | 13647S |
| STING | Ms monoclonal | WB | R&D | MAB7169 |
| pSTING (Ser366) (D7C3S) | Rbt monoclonal | WB | CST | 19781S |
| pSTING (Ser366) (D8K6H) | Rbt monoclonal | IF | CST | 40818S |
| TBK1 (E8I3G) | Rbt monoclonal | WB | CST | 38066S |
| **TBK1 (E9H5S)** | Ms monoclonal | WB | CST | 51872S |
| **pTBK1 (Ser172) (D52C2)** | Rbt monoclonal | WB | CST | 5483S |
| **IRF3 (D6I4C)** | Rbt monoclonal | WB | CST | 11904S |
| **pIRF3 (Ser386) (E7J8G)** | Rbt monoclonal | WB | CST | 37829S |
| **STAT1** | Rbt monoclonal | WB | CST | 9172S |
| **cGAS (D1D3G)** | Rbt monoclonal | WB | CST | 15102S |
| **pSTAT1 (Tyr 701) (58D6)** | Rbt monoclonal | WB | CST | 9167S |
| **MX1 (D3W7I)** | Rbt monoclonal | WB | CST | 37849S |
| **HRS (D7T5N)** | Rbt monoclonal | WB and IF | CST | 15087S |
| **LC3B** | Rbt monoclonal | WB and IF | CST | 2775S |
| **p62 (D5L7G)** | Rbt monoclonal | IF | CST | 88588S |
| **Ubc13** | Rbt monoclonal | WB | CST | 4919 |
| **CD63 (EPR22458-280)** | Rbt monoclonal | IF | Abcam | ab252919 |
| **CD63 (MX-49.129.5)** | Ms monoclonal | IF | Santa Cruz | sc-5275 |
| **VPS37A (G-3)** | Ms monoclonal | WB | Santa Cruz | sc-376978 |
| **NBR1 (4BR)** | Ms monoclonal | IF | Santa Cruz | sc-130380 |
| **UBAP1** | Rbt polyclonal | WB | Proteintech | 12385-1-AP |
| **RFP (6G6)** | Ms monoclonal | WB | Chromotek | 6g6-100 |
| **ATG5 (D5F5U)** | Rbt monoclonal | WB | CST | 12994S |
| **ATG16L1 (D6D5)** | Rbt monoclonal | WB | CST | 8089S |
| **GM130 (35/GM130)** | Ms monoclonal | IF | BD Biosciences | 610823 |
| **EEA1 (14/EEA1)** | Ms monoclonal | IF | BD Biosciences | 610457 |
| **β-actin (AC-15)** | Ms monoclonal | WB | Abcam | ab6276 |
| **α-Tubulin (DM1A)** | Ms monoclonal | WB | Thermo Fisher | 14-4502-82 |
| **Vinculin** | Rbt monoclonal | WB | CST | 4650S |
| **HRP Anti-beta Actin antibody [AC-15]** | Ms monoclonal | WB | Abcam | ab49900 |
| **HRP-anti β-actin (mAbcam 8226)** | Ms Monoclonal - HRP conjugate | WB | Abcam | ab20272 |
| **HRP-Streptavidin** | NA | WB | BioLegend | 405210 |
| **HA** | Rat monoclonal | IF | Millipore Sigma | 11867423001 |
| **DYKDDDDK Tag (D6W5B) - FLAG** | Rabbit monoclonal | WB | CST | 14793S |
| **Cy5-Streptavidin** | NA | IF | Biolegend | 405209 |
| **Alexa Fluor 647 anti-HA.11** | Biolegend | FC | Biolegend | 682404 |
| **Alexa 488 anti-mouse IgG (H+L)** | Gt polyclonal | IF | Thermo Fisher | A-11029 |
| **Alexa 555 F(ab')<sub>2</sub> anti-Rabbit IgG** | Gt polyclonal | IF | Thermo Fisher | A-21430 |
| **Alexa 647 anti-rat IgG (H+L)** | Gt polyclonal | IF | Thermo Fisher | A-21247 |
| **Peroxidase AffiniPure Goat Anti-Rabbit IgG (H+L)** | Gt polyclonal | WB | Jackson ImmunoResearch | 111-035-144 |
| **Peroxidase AffiniPure Goat Anti-Mouse IgG (H+L)** | Gt polyclonal | WB | Jackson ImmunoResearch | 111-035-146 |

Rbt: rabbit; Ms: mouse; Gt: goat.
WB: western blot; IF: immunofluorescence; FC: flow cytometry.

## RNA sequencing reagents and conditions

| RT mix 1 |  |  |
| --- | --- | --- |
| Reagent | Reaction conc. | μL required |
| PEG 8000 (50% solution) | 5% | 3.2 |
| Triton X-100 (10% solution) | 0.10% | 0.24 |
| RNAse Inhibitor (40μ/μL) | 0.5μ/μL | 0.32 |
| Smartseq3_OligodT30VN(100μM) | 0.5μM | 0.16 |
| dNTPs (10mM/each) | 0.5mM/each | 1.6 |
| mRNA template |  | μL for 15ng |
| Nuclease Free Water |  | Up to 24μL |
| Total uL |  | 24 |

| RT mix 2 |  |  |
| --- | --- | --- |
| Reagent | Reaction conc. | μL required |
| Tris-HCl pH 8.5 (1M) | 25mM | 0.8 |
| NaCl (1M) | 30mM | 0.96 |
| MgCl<sub>2</sub> (100mM) | 2.5mM | 0.8 |
| GTP (100mM) | 1mM | 0.32 |
| DTT (100mM) | 8mM | 2.56 |
| RNAse Inhibitor (40u/uL) | 0.5u/uL | 0.4 |
| TSO (100uM) | 2uM | 0.64 |
| Maxima H-minus RT enzyme (200U/uL) | 2u/uL | 0.32 |
| Nuclease Free Water |  | 1.2 |
| Total uL |  | 8 |

| First-strand synthesis |  |  |
| --- | --- | --- |
| Temperature | Time | Cycles |
| 42 °C | 90 min | 1x |
| 50 °C | 2 min | 10x |
| 42 °C | 2 min |  |
| 85 °C | 5 min | 1x |

| cDNA PCR mix |  |  |
| --- | --- | --- |
| Reagent | Reaction conc. | uL per. reaction |
| Kapa HiFi HotStart buffer (5X) | 1X | 10 |
| dNTPs (10mM/each) | 0.3mM/each | 1.5 |
| MgCl<sub>2</sub> (100mM) | 0.5mM | 0.25 |
| Fwd Primer (100uM) AAO270 | 0.5uM | 0.25 |
| Rev Primer (10uM) AAO271 | 0.1uM | 0.5 |
| First-strand synthesis reaction |  | 20 |
| Polymerase (1U/uL) | 0.02U/uL | 1 |
| Nuclease Free Water |  | 16.5 |
| Total uL |  | 30 |

| cDNA amplification |  |  |
| --- | --- | --- |
| Temperature | Time | Cycles |
| 98 °C | 3 min | 1x |
| 98 °C | 20 sec | 25x |
| 65°C | 30 sec |  |
| 72 °C | 6 min |  |
| 72 °C | 5 min | 1x |
| 4 °C | Hold |  |

| Tagmentation mix |  |  |
| --- | --- | --- |
| Reagent | Reaction conc. | uL per. reaction |
| Tagmentation buffer (4x)<br>(Tris-HCl pH 7.5 - 40mM, MgCl<sub>2</sub> - 20mM, DMF- 20%) | 1X | 2 |
| Amplicon Tagmentation Mix (Tn5) |  | 0.32 |
| UltraPure water |  | 1.68 |
| Total |  | 4 |

| Final library amplification mix |  |  |
| --- | --- | --- |
| Reagent | Reaction conc. | uL per. reaction |
| Phusion HF buffer (5X) | 1X | 5.6 |
| dNTPs (10mM/each) | 0.2mM/each | 0.6 |
| Phusion HF (2U/uL) | 0.01U/uL | 0.16 |
| Nextera Index Primers (0.5µM) | 0.1µM | 6 |
| H2O |  | 5.64 |
| Tagmented cDNA |  | 10 |
| Total uL |  | 28 |

| Final library amplification |  |  |
| --- | --- | --- |
| Temperature | Time | Cycles |
| 72 °C | 3 min | 1x |
| 98 °C | 3 min | 1x |
| 98 °C | 10 sec | 12x |
| 55 °C | 30 sec |  |
| 72 °C | 30 sec |  |
| 72 °C | 5 min | 1x |
| 4 °C | Hold |  |

---

**Figure S1. Related to Figure 1.**
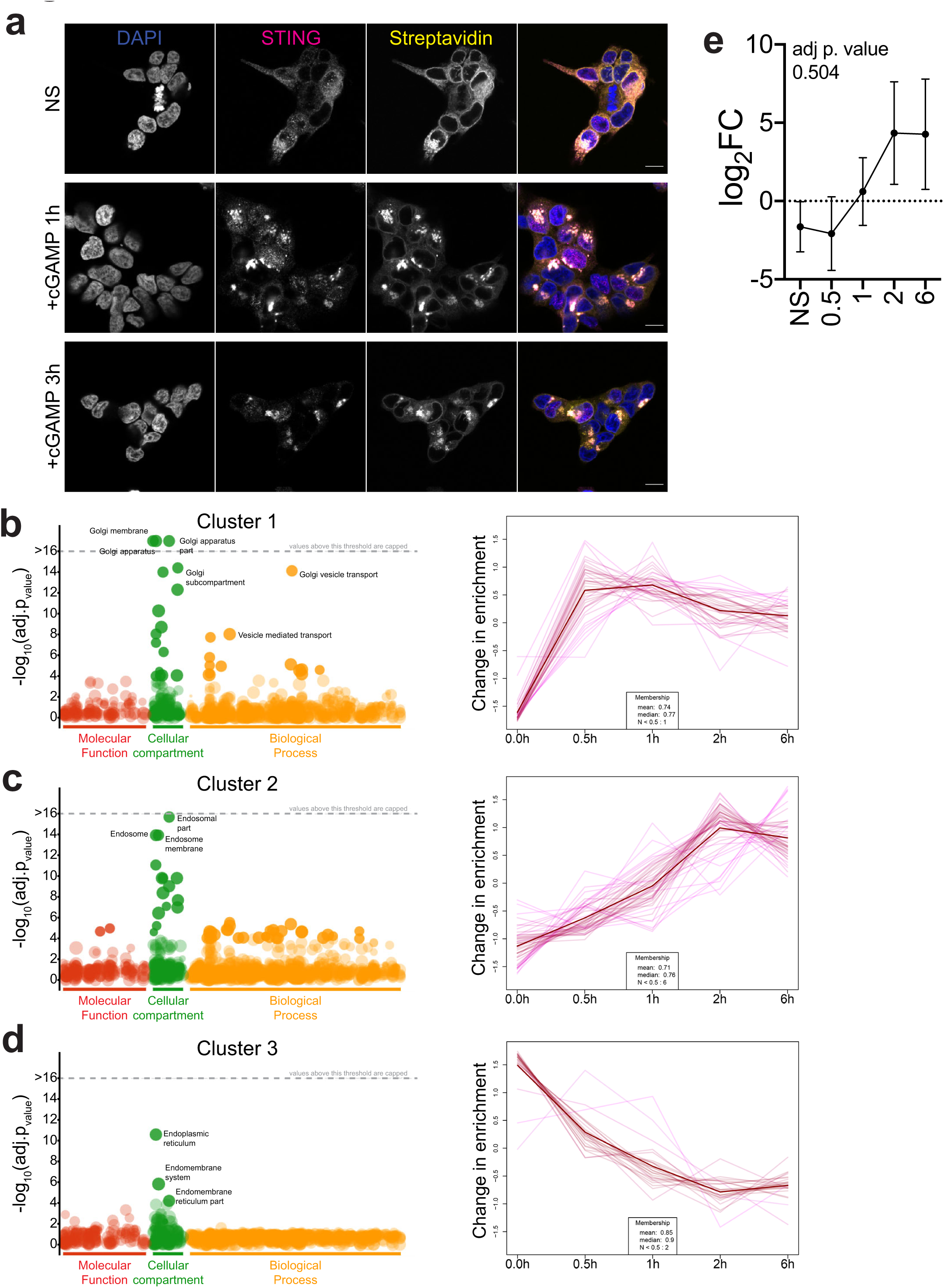
**a)** Immunofluorescence of DAPI (blue) STING-TurboID (magenta) and Streptavidin-Cy5 (yellow) at the indicated time-points in 293T STING-TurboID cells post cGAMP stimulation or non-stimulated (NS). One representative field of n≥3 fields. Scale bar is 10µm. **b)** Cluster 1 from cluster analysis. Molecular Function, Cellular compartment and Biological process GO terms enrichment (left) and plot of change in enrichment of statistically significant proteins at the indicated time-points. **c)** Same as in b) for Cluster 2. **d)** Same as in b) for Cluster 3. **e)** log2FC of p62 at the indicated time-points.

**Figure S2. Related to Figure 1.**
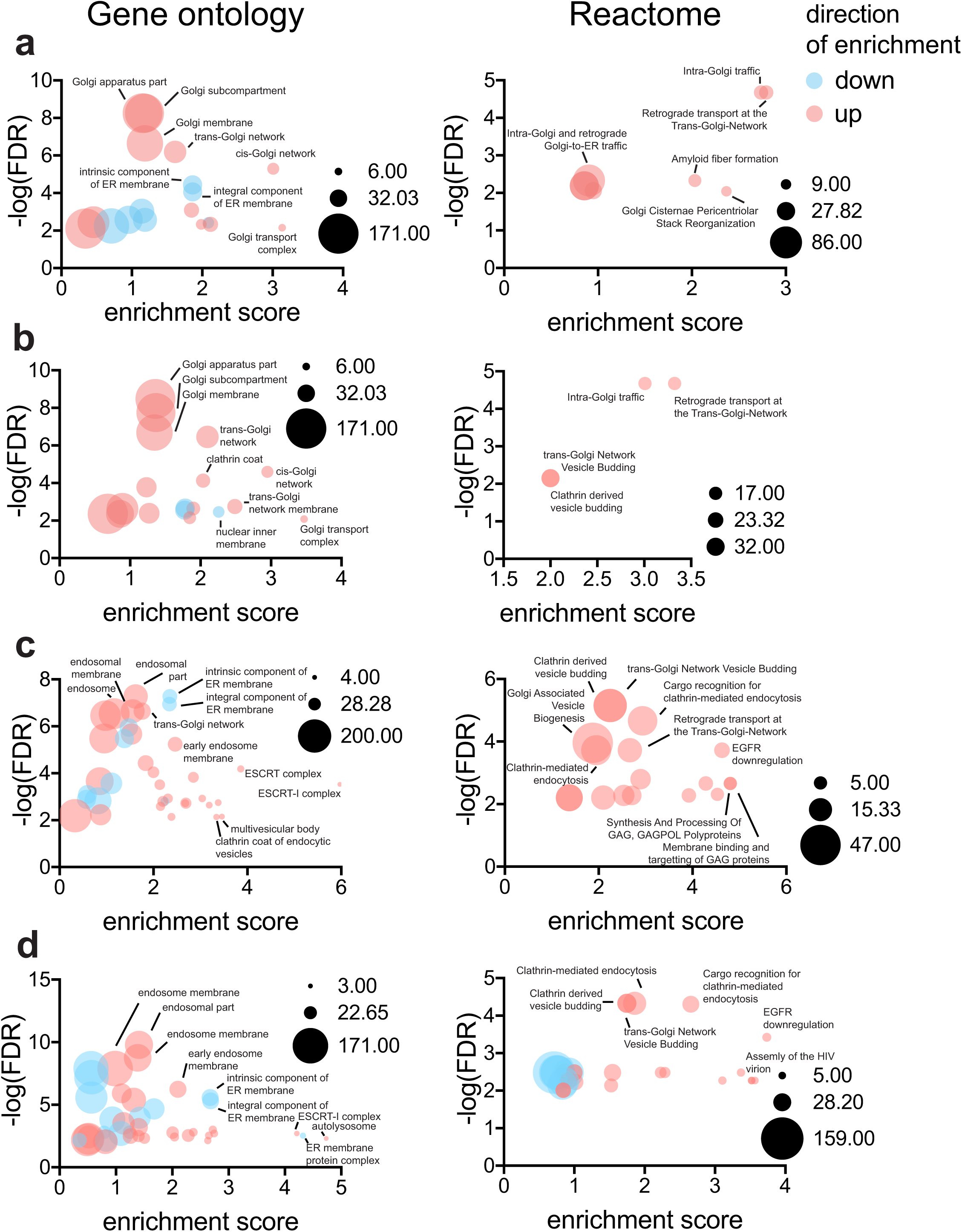
Gene Ontology (GO) (left panels) and Reactome (right panels) enrichments calculated through STRING for the full TurboID dataset at **a)** 30 minutes, **b)** 1 hour, **c)** 2 hours and **d)** 6 hours post cGAMP stimulation. Terms that are positively enriched are in red, terms that are negatively enriched are in blue. Size of bubbles represent the number of genes mapped in each category. FDR: False Discovery Rate.

**Figure S3. Related to Figure 2.**
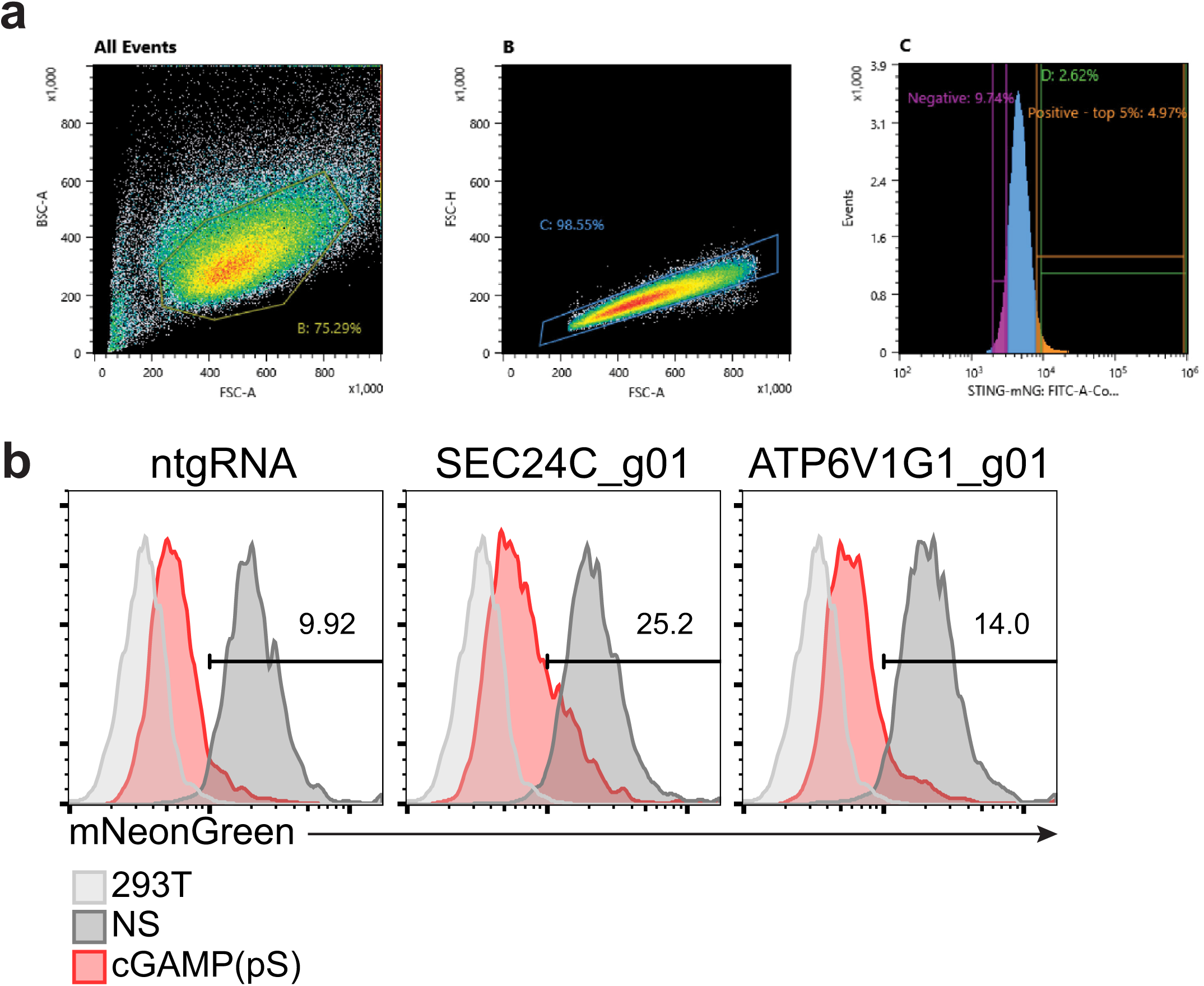
**a)** Gating strategy for the genome-wide CRISPR screen. Gate-D in Panel C not used. **b)** mNeonGreen levels in reporter spCas9 expressing 293T STING-mNeonGreen cell lines transduced with the indicated sgRNAs before (dark gray – NS) or after (red) stimulation with 1µg/ml 2’3’-cGAMP(pS)2 for 24 hours. Line represents gating strategy, and numbers represent %STING-mNeonGreen positive cells post stimulation. 293T (light gray) are shown as a reference for mNeonGreen negative cells. One representative plot of n=2 independent experiments with n=2 technical replicates per experiment.

**Figure S4. Related to Figure 3.**
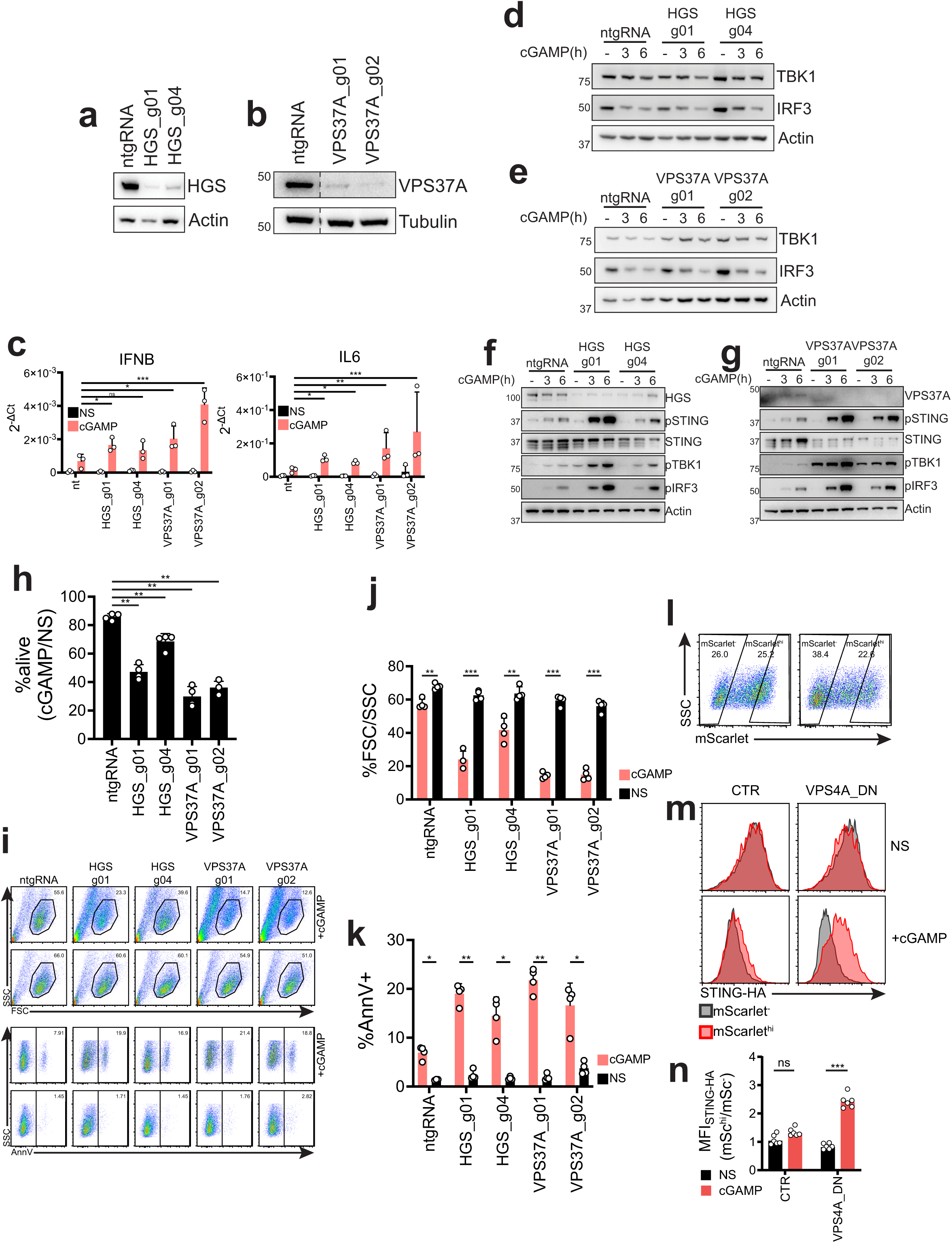
**a)** HGS KO efficiency relative to Fig. 3b, 3c. **b)** VPS37A KO efficiency relative to Fig. 3b, 3c. **c)** 2^-ΔCt^ values related to Fig. 3j. n=3 independent experiments. One-way ANOVA on log-transformed data with Dunnet multiple comparison test. *p<0.05, **p<0.01, ***p<0.001, ns=not significant. **d)** Immunoblot of the indicated proteins in BJ1 KO for HGS, relative to 3h. **e)** Same as in d) for VPS37A, relative to Fig. 3i. **f)** Immunoblot of the indicated proteins in U937 KO for HGS with two independent guides per gene or transduced with a non-targeting sgRNA (CTR). 20µg/ml cGAMP stimulation (in medium) was for 6 hours. One representative blot of n=3 independent experiments. **g)** Same as in f), for VPS37A. **h)** U937 cells were treated with 20µg/ml cGAMP (in medium) for 24 hours and alive cells were quantified with Cell Titer Glo. Alive cells are shown as percentage ratio RLU in cGAMP treated samples over non-stimulated samples. n=4 independent experiments. One-Way ANOVA with Dunnet multiple comparison test **i)** FSC/SSC and Annexin V plots for U937 treated as in g). One representative experiment of n=4 independent experiments. **j)** Percentage of cells in FSC/SSC gate for cells treated as in i). n=4 independent experiments. One-Way ANOVA with Holm-Sidak multiple comparison test. **k)** Percentage Annexin V positive cells in FSC/SSC gate for cells treated as in i). n=4 independent experiments. One-Way ANOVA with Holm-Sidak multiple comparison test. ****p<0.0001, ***p<0.001, **p<0.01, *p<0.05. **l)** Expression of mScarlet-VPS4ADN (VPS4A E228Q) after transfection in 293T stably expressing STING-HA and gating strategy for mScarlet^-^ and mScarlet^hi^ cells. **m)** STING-HA levels in cells as in l) in mScarlet^-^ (dark grey) and mScarlet^hi^ (red) populations that were either non-stimulated (NS) or treated with 2µg/ml cGAMP (in perm buffer) for 6 hours. One representative plot of n=3 independent experiments with n=2 technical replicates per experiment. **n)** MFI of STING-HA signals shown as a ratio of MFI of the mScarlet^hi^ population over the MFI of the mScarlet^-^ population. n=3 independent experiments with n=2 technical replicates per experiment. Each dot represents an individual replicate. One-way ANOVA with Dunnet multiple comparisons test. ***p<0.001, ns: not-significant

**Figure S5. Related to Figure 4.**
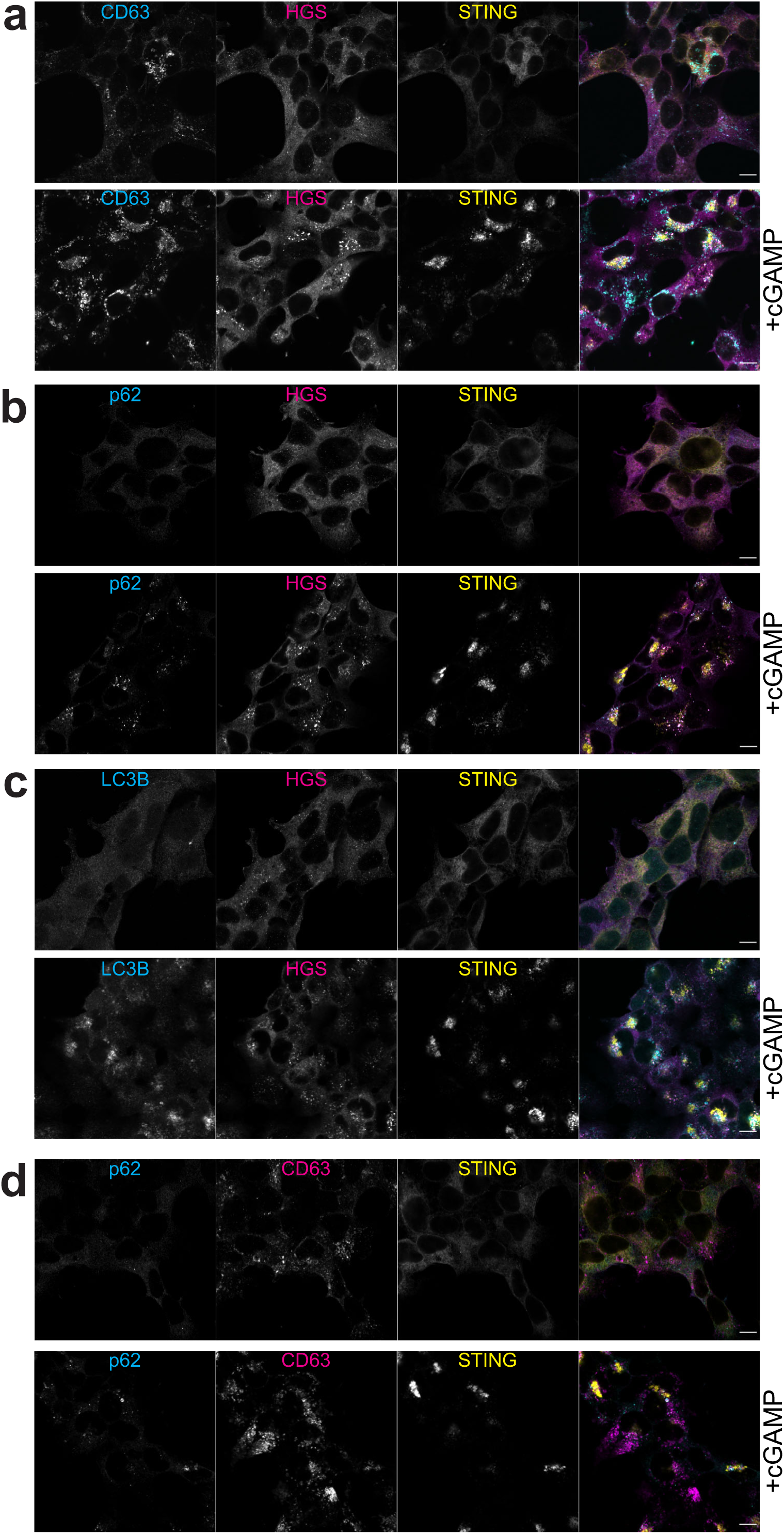
Single color and merge of immunofluorescence of **a)** CD63 (cyan), HGS (magenta) and STING (yellow), **b)** p62 (cyan), HGS (magenta) and STING (yellow), **c)** LC3B (cyan), HGS (magenta) and STING (yellow), **d)** p62 (cyan), CD63 (magenta) and STING (yellow) in 293T stably expressing STING-HA non-stimulated or stimulated with cGAMP relative to Figure 4a-d. Scale bar is 10µm. One representative field of n≥3 independent fields in n=3 independent experiments.

**Figure S6. Related to Figure 5.**
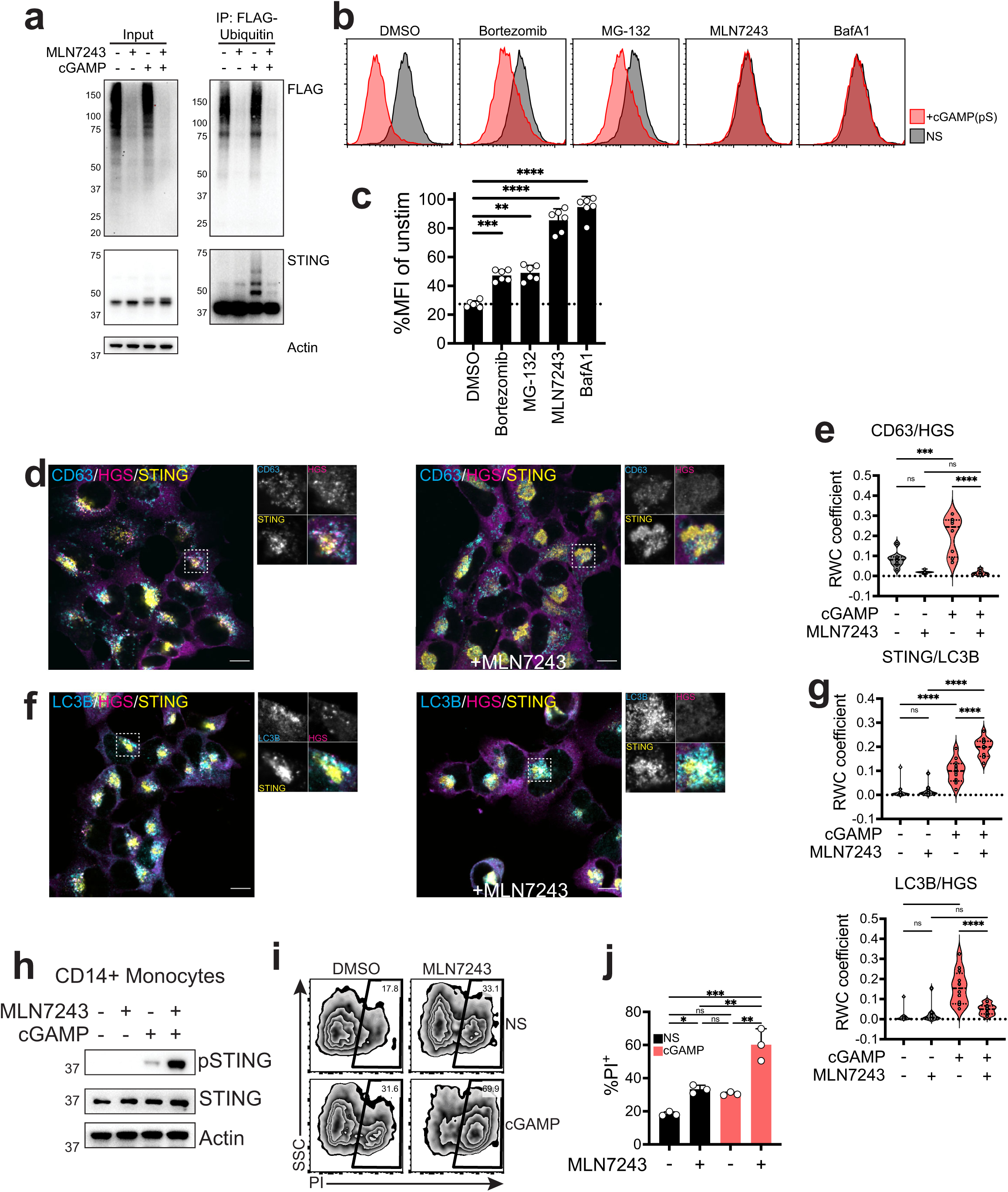
**a)** Immunoblot of the indicated proteins in 293T stably expressing STING-HA and FLAG-ubiquitin in the input or after FLAG pulldown (IP: Flag-Ubiquitin) for cells stimulated with 2µg/ml cGAMP (in perm buffer) in presence or absence of 0.5µM MLN7243 for 2 hours. One representative blot of n=2 independent experiments. **b)** STING-mNG levels in 293T STING-mNG cells treated with 4µg/ml cGAMP(pS)2 (in medium) for 6 hours in presence of the indicated drugs. One representative experiment of n=3 independent experiments with n=2 technical replicates. **c)** Percentage MFI of STING-mNG signal expressed as ratio MFI of stimulated cells as in b) over MFI of unstimulated cells. n=3 independent experiments with n=2 technical replicates. One-way ANOVA with Dunnet multiple comparisons test. ****p<0.0001, ***p<0.001, **p<0.01. **d)** Immunofluorescence of CD63 (cyan), HGS (magenta) and STING (yellow) in absence (left) or in presence (right) of MLN7243 in 293T stably expressing STING-HA after stimulation with 2µg/ml cGAMP (in perm buffer) for 2 hours. Dashed boxes represent the cropped regions shown in the right panels. One representative field of n≥5 fields in n=2 independent experiments. Scale bar is 10µm. Control non-stimulated cells are in Fig. S7. **e)** Rank Weighted Colocalization (RWC) coefficient for CD63 colocalization in HGS foci in cells stimulated as in d). Each dot represents colocalization calculated in a field. n=2 independent experiments with n≥5 fields. One way ANOVA with post-hoc Tukey test. *p<0.05, **p<0.01, ***p<0.001, ****p<0.0001, ns=not significant. **f)** Immunofluorescence of CD63 (cyan), HGS (magenta) and STING (yellow) in absence (left) or in presence (right) of MLN7243 in cells treated as in d). Dashed boxes represent the cropped regions shown in the right panels. One representative field of n≥5 fields in n=2 independent experiments. Scale bar is 10µm. Control non-stimulated cells are in Fig. S7. **g)** RWC for LC3B colocalization in STING foci (top) or LC3B colocalization in HGS foci (bottom) in cells stimulated as in f). Each dot represents colocalization calculated in a field. n=2 independent experiments with n≥5 fields. One way ANOVA with post-hoc Tukey test. *p<0.05, **p<0.01, ***p<0.001, ****p<0.0001, ns=not significant. **h)** Immunoblot of the indicated proteins in primary CD14+ monocytes treated with 5µg/ml cGAMP (in medium) for 8h. One donor representative of n=3 independent donors. **i)** Propidium Iodide (PI) staining in CD14+ monocytes treated as in h). One representative donor of n=3 independent donors. **j)** %PI^+^ CD14+ monocytes treated as in h. Each dot represents an independent donor (n=3). One-way ANOVA with post-hoc Tukey test. *p<0.05, **p<0.01, ***p<0.001, ns=not significant.

**Figure S7. Related to Figure 5 and Figure S6.**
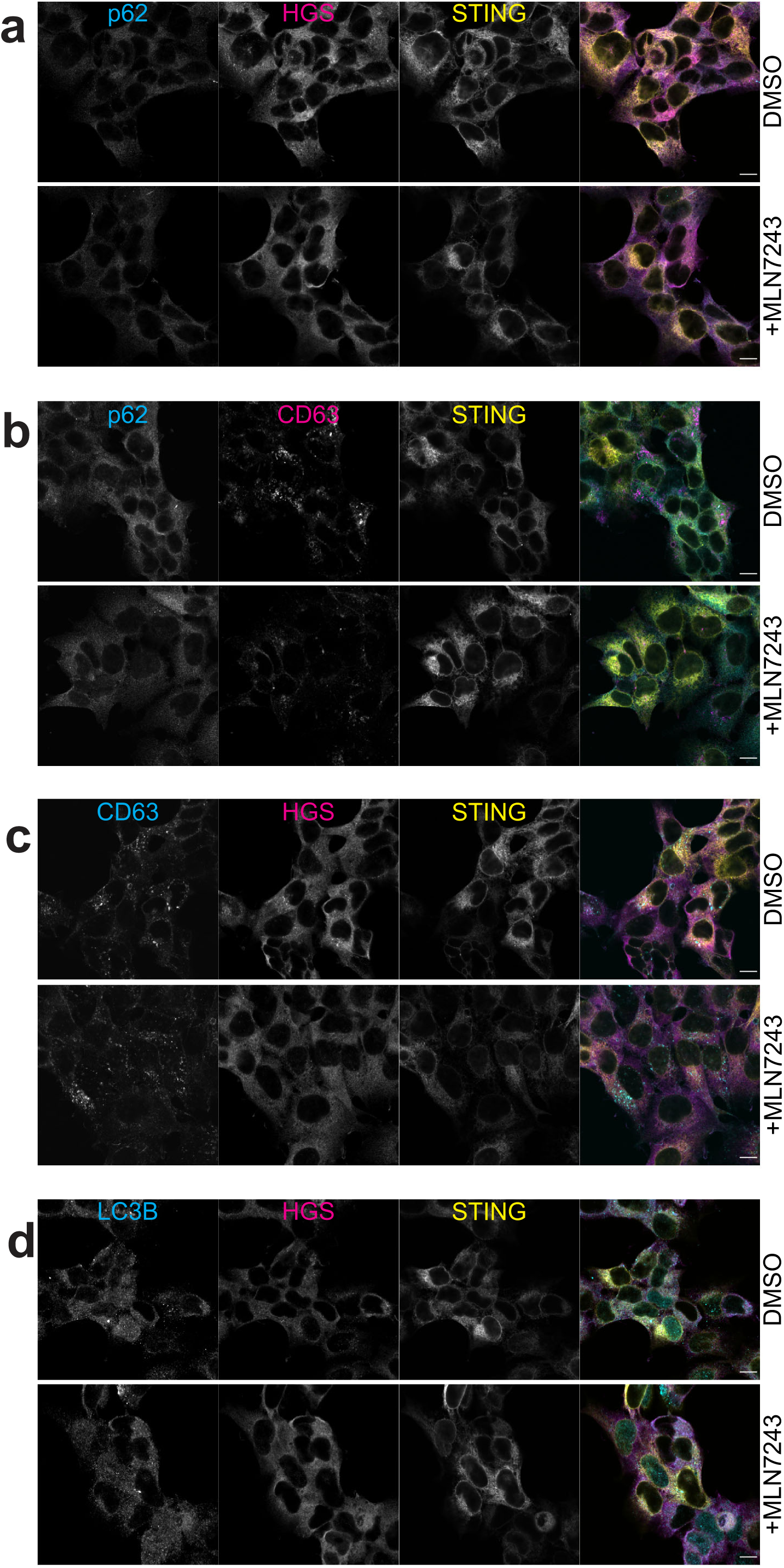
Immunofluorescence of **a)** p62 (cyan), HGS (magenta) and STING (yellow), **b)** p62 (cyan), CD63 (magenta) and STING (yellow), **c)** CD63 (cyan), HGS (magenta) and STING (yellow), **d)** LC3B (cyan), CD63 (magenta) and STING (yellow) in 293T stably expressing STING-HA treated (bottom panels) or non-treated (DMSO – top panels) with MLN7243.

**Figure S8. Related to Figure 6.**
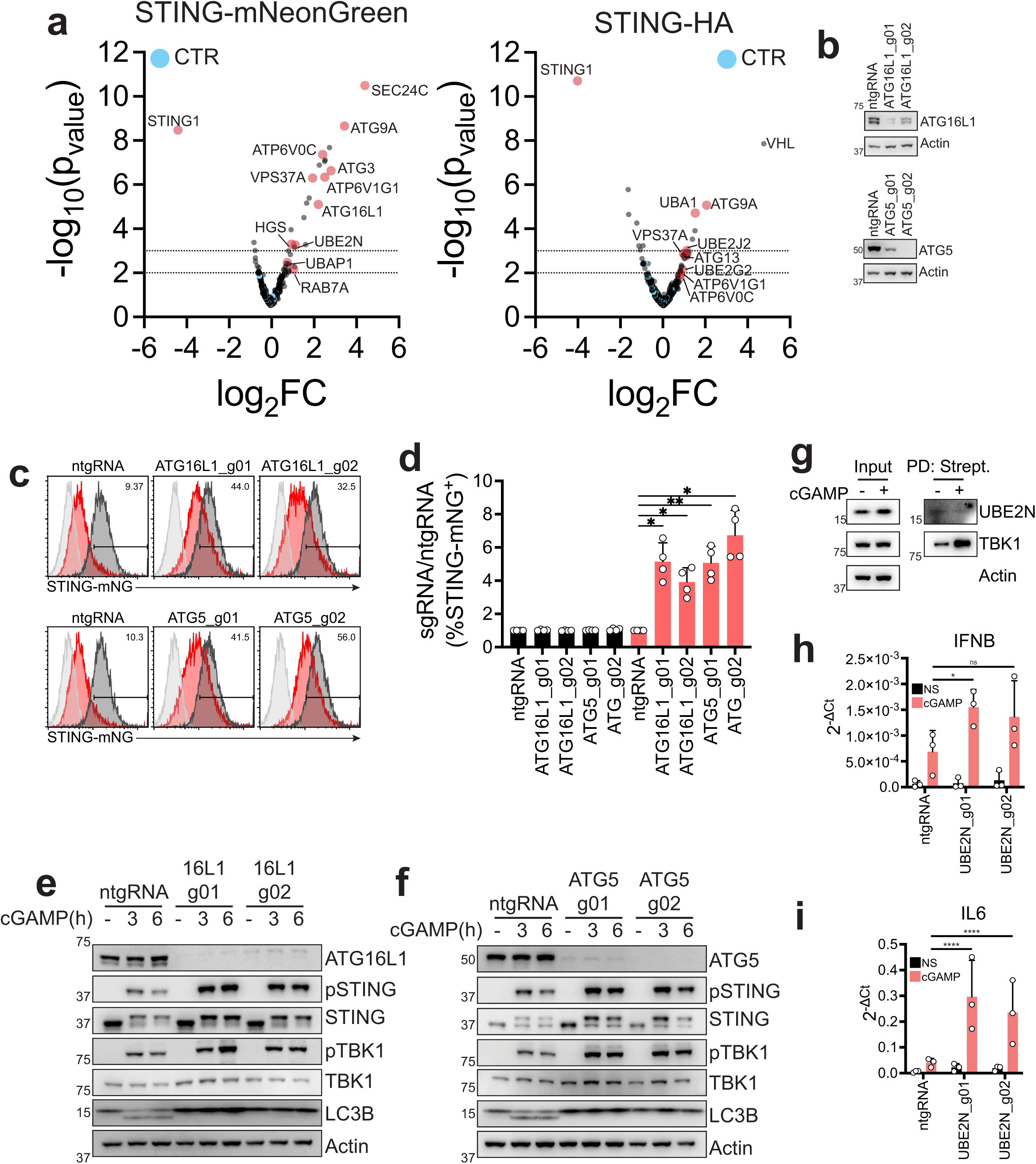
**a)** Volcano-plos of log_2_ fold change (log_2_FC) vs -log_10_(p_value_) after sequencing and analysis of the targeted screens in STING-mNeonGreen and STING-HA cell lines. VHL scoring highly in the STING-HA screen is due to the use of a hPGK promoter to drive STING-HA expression. KO of VHL stabilizes HIF1ɑ which drives transcription from the HRE elements present in the hPGK promoter. **b)** Immunoblot of the indicated proteins in 293T STING-mNG KO for the indicated genes. **c)** mNeonGreen levels in 293T STING-mNeonGreen cell lines same as in b) before (dark gray – NS) or after (red) stimulation with 4µg/ml 2’3’-cGAMP(pS)2 (in medium) for 6 hours. Line represents gating strategy and numbers represent %STING-mNG positive cells post stimulation. 293T (light gray) are shown as a reference for mNG negative cells. One representative plot of n=2 independent experiments with n=2 technical replicates per experiment. **d)** Percentage of STING-mNeonGreen (mNG) positive cells in cells stimulated as in c). Shown is ratio %STING-mNG positive of each sgRNA over %STING-mNG positive cells of the control non-targeting sgRNA (ntgRNA). n=2 independent experiments with n=2 technical replicates per experiment. Each dot represents an individual replicate. One-way ANOVA with Dunnet multiple comparison test. *p<0.05, **p<0.01. **e)** Immunoblot of the indicated proteins in BJ1 fibroblasts KO with two independent guides for ATG16L1. One representative experiment of n=2 independent experiments. **f)** Immunoblot of the indicated proteins in BJ1 fibroblasts KO with two independent guides for ATG5. One representative experiment of n=2 independent experiments. **g)** Immunoblot of the indicated proteins in 293T STING-TurboID stimulated for 1 hour with 2µg/ml cGAMP. **h)** 2-ΔCt values related to Fig. 6f **i)** and Fig. 6g. n=3 independent experiments. One-way ANOVA on log-transformed data with Dunnet multiple comparison test. *p<0.05, ****p<0.0001, ns=not significant.

**Figure S9. Related to Figure 7.**
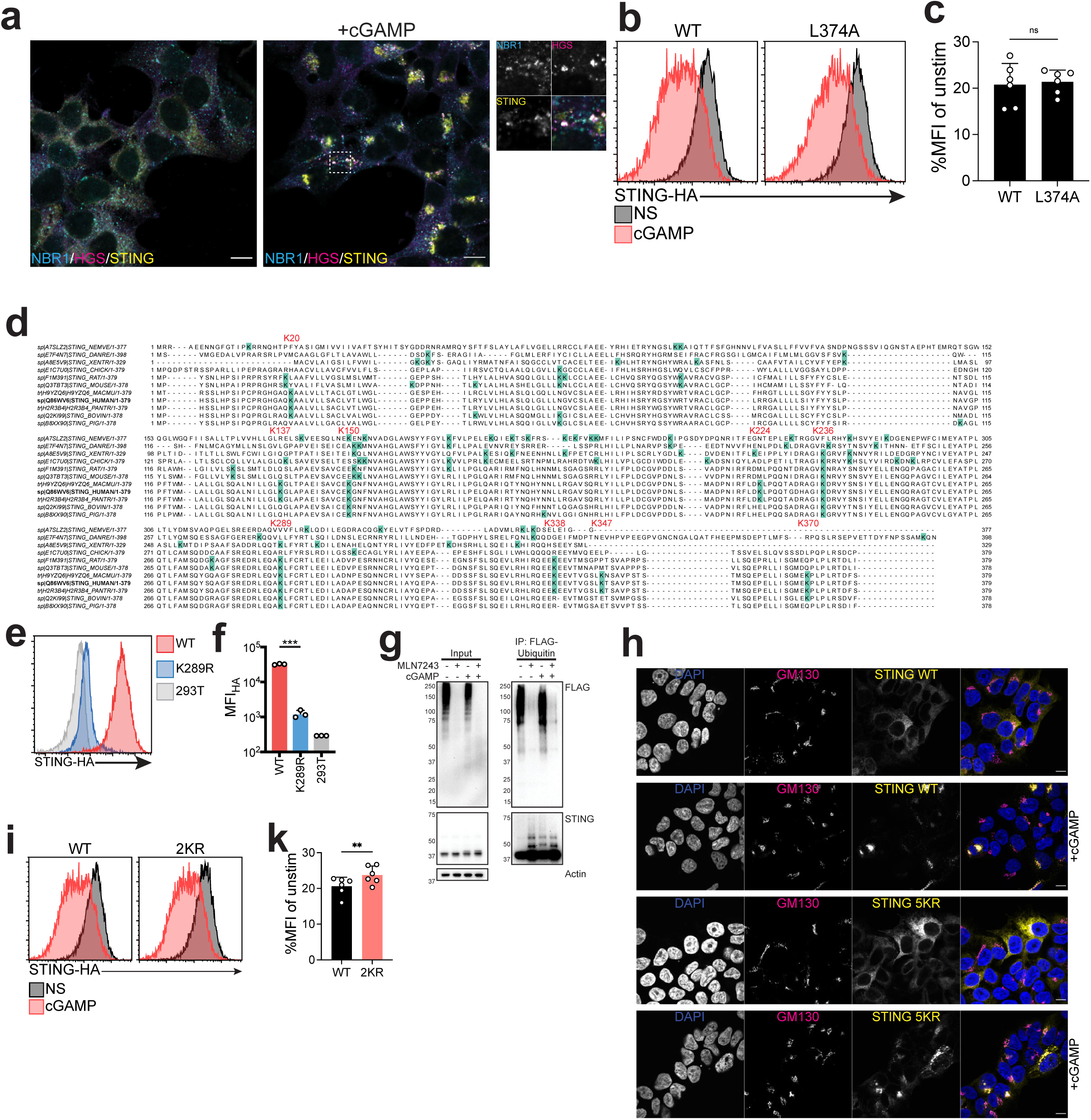
**a)** Immunofluorescence of NBR1 (cyan), HGS (magenta) and STING (yellow) in 293T STING-HA cells stimulated with 2µg/ml cGAMP (in perm buffer) for 2h. Dashed boxes represent the cropped regions shown in the right panels. One representative field of n≥5 fields in n=2 independent experiments. Scale bar is 10µm. **b)** HA level in 293T cells expressing STING WT-HA or STING L374A-HA stimulated with 2µg/ml cGAMP (in perm buffer) for 6h. One experiment representative of n=3 independent experiments with n=2 technical replicates. **c)** Median Fluorescence Intensity (MFI) of cells as in b) shown as %MFI of cGAMP stimulated over non-stimulated (NS) for each mutant. n=3 independent experiments with n=2 technical replicates per experiment. Each dot represents an individual replicate. Paired t-test. ns=not significant. **d)** Alignment of STING in different species. Lysines are highlighted and red numbers refer to positions in human STING. **e)** HA levels in 293T stably expressing HA labeled STING WT or K289R. One representative plot of n=3 technical replicates. **f)** Median Fluorescence intensity for the STING mutants as in e). Each dot represents one technical replicate. Paired t-test, ***p<0.001. **g)** Immunoblot of the indicated proteins in 293T stably expressing STING 5KR-HA and FLAG-ubiquitin in the input or after FLAG pulldown (IP: Flag-Ubiquitin) for cells stimulated with 2µg/ml cGAMP (in perm buffer) in presence or absence of MLN7243 for 2 hours. One representative blot of n=2 independent experiments. **h)** Immunofluorescence of DAPI (blue), GM130 (magenta) and STING (yellow) in 293T stably expressing either STING WT or STING 5KR stimulated with 2µg/ml cGAMP (in perm buffer) for 2h. **i)** HA levels in 293T stably expressing STING WT or STING 2KR (K338R/K370R) stimulated with 2µg/ml cGAMP (in perm buffer) for 6h. One representative experiment of n=3 independent experiments with n=2 technical replicates. **k)** Median Fluorescence Intensity (MFI) of cells as in i) shown as %MFI of cGAMP stimulated over non-stimulated (NS) for each mutant. n=3 independent experiments with n=2 technical replicates per experiment. Each dot represents an individual replicate. Paired t-test. **p<0.01.

**Figure S10. Related to Figure 8.**
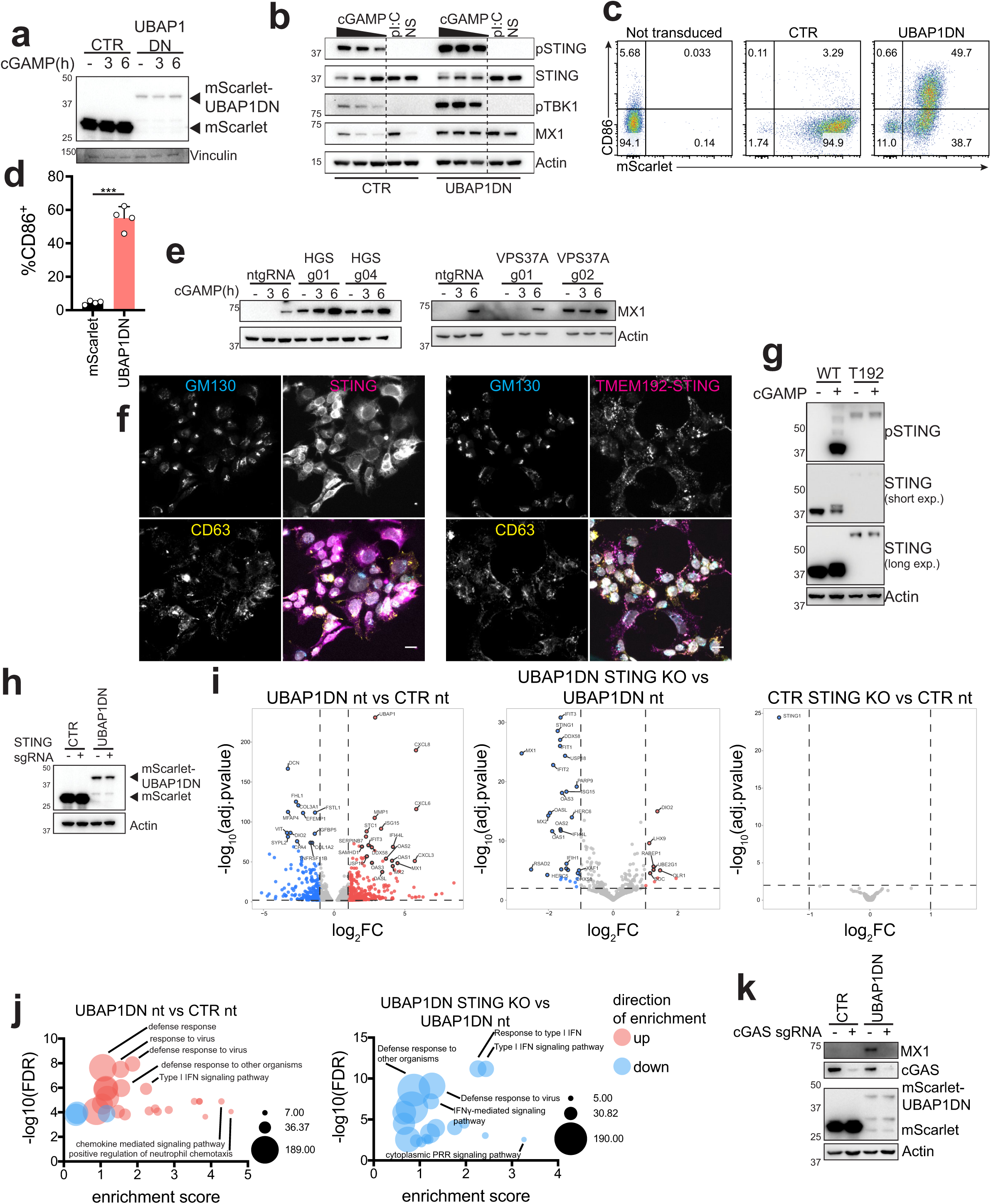
**a)** Immunoblot of the indicated proteins. Related to Fig. 7f. **b)** Immunoblot of the indicated proteins in primary human MDDCs transduced with either a control vector (mScarlet) or mScarlet-UBAP1DN stimulated with cGAMP (top dose 10µg/ml, 1:2 dilutions) or pI:C+Lipofectamine for 6h. One representative donor of n=4 donors in n=2 independent experiments. **c)** CD86 and mScarlet expression in cells transduced as in b). One representative donor of n=4 donors in n=2 independent experiments. **d)** Quantification of %CD86+ cells in mScarlet+ gate. n=4 donors in n=2 independent experiments. Paired t-test. ***p<0.001. **e)** Immunoblot of the indicated proteins in BJ1 fibroblasts non-stimulated (-) or stimulated with cGAMP for the indicated times. One representative experiment of n=3 experiments. **f)** Immunofluorescence of GM130 (cyan), HA (magenta) and CD63 (yellow) in 293T stably expressing STING-HA (left panel) or TMEM192-STING-HA (right panel). DAPI is in grey in the merged image. Scale bar is 20µm. One field representative of n≥5 fields per construct of n=2 technical replicates. **g)** Immunoblot of the indicated proteins in 293T stably expressing WT STING-HA (WT) or TMEM192-STING-HA (T192) stimulated or not with 2µg/ml cGAMP for 2h. One experiment representative of n=2 independent experiments with n=2 technical replicates. **h** Immunoblot of the indicated proteins. Related to Fig. 7g. **i)** Volcano-plots of the differentially expressed genes (DEGs) for the indicated comparisons. Dashed lines to indicate significant DEGs are drawn at log_2_FC ≥1 and log_2_FC≤-1 and -log_10_(adjusted p value)≥2. Significant downregulated genes are in blue, upregulated in red. nt: non-targeting sgRNA **j)** GO analysis of significantly enriched processes in the indicated conditions. Positively enriched terms are in red, negatively enriched in blue. Size of bubbles represents the number of mapped genes in each category. FDR: False Discovery Rate. **k)** Immunoblot of the indicated proteins in BJ1 expressing mScarlet (CTR) or mScarlet-UBAP1DN transduced with spCas9 and a control sgRNA or a cGAS targeting sgRNA. One experiment representative of n=3 independent experiments.

**Figure S11.**
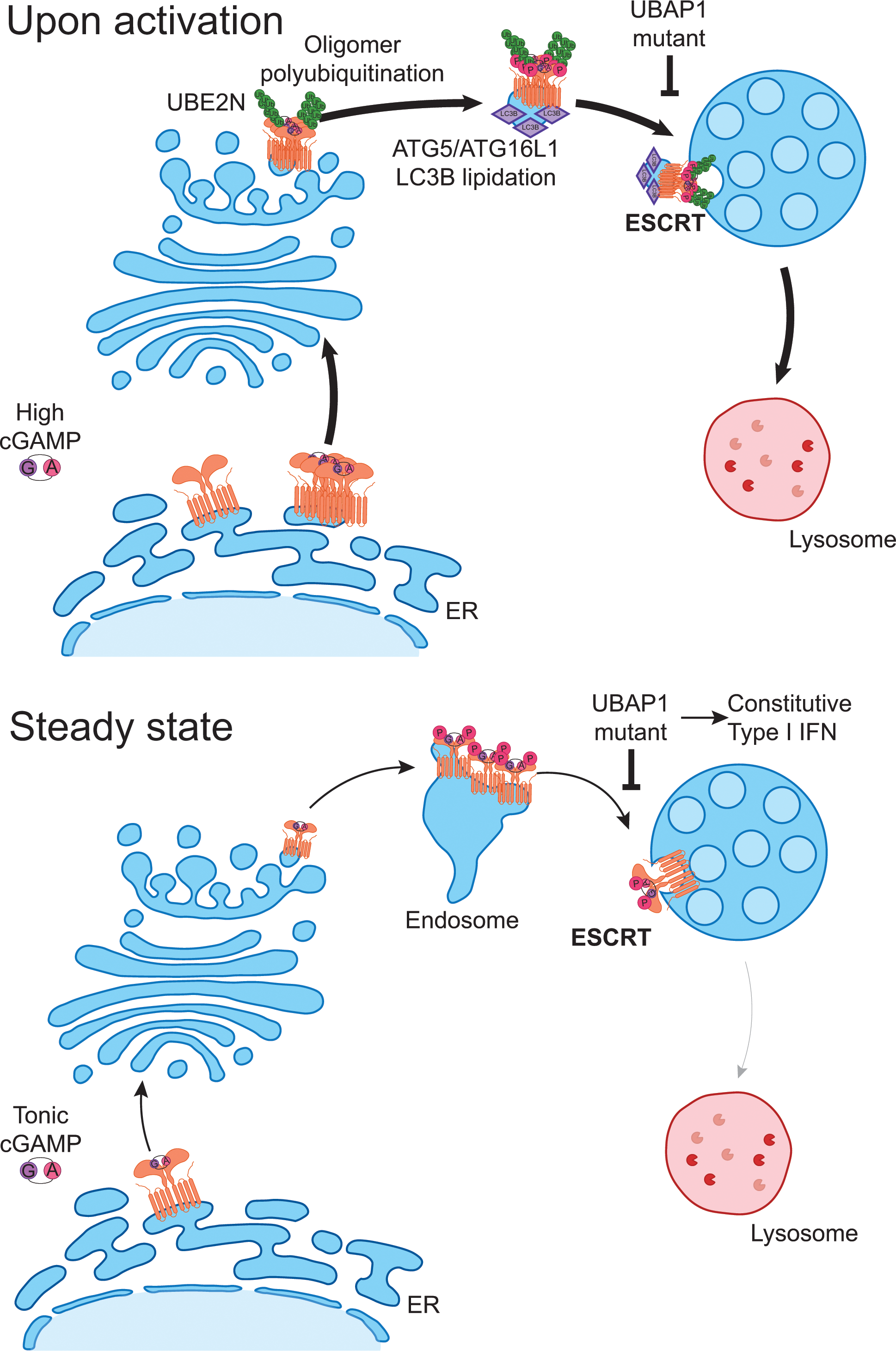
ESCRT-dependent STING degradation curtails steady-state and cGAMP-induced signaling. *Top.* Activated STING traffics from the ER to the Golgi and then to the endosome. High levels of intracellular cGAMP drive STING oligomerization which in turns lead to UBE2N dependent STING polyubiquitination and ATG5/ATG16L1 dependent LC3B lipidation of STING containing vesicles. Ubiquitination of STING drives its association with ESCRT at the late endosome. Association of ESCRT to STING creates an organizing center for fusion with the endolysosomal compartment leading to STING degradation. Pathogenic mutants of the ESCRT-I subunit UBAP1 block this process and lead to exacerbated STING responses. *Bottom.* At steady state, cGAS primes tonic STING trafficking between the ER and the lysosomes through the endosomal compartment. ESCRT ensures removal of STING at steady state preventing spontaneous activation of the sensor. A UBAP1 mutant, or KO of HGS and VPS37A, blocking ESCRT function leads to accumulation of phosphorylated STING at the endosome consequently driving constitutive STING activation. Mutations in genes regulating post-Golgi STING trafficking could therefore lead to spontaneous activation of the sensor and underlie disease.

